# Representational integration and differentiation in the human hippocampus following goal-directed navigation

**DOI:** 10.1101/2022.04.12.488078

**Authors:** Corey Fernandez, Jiefeng Jiang, Shao-Fang Wang, Hannah L. Choi, Anthony D. Wagner

## Abstract

As we learn, dynamic memory processes build structured knowledge across our experiences. Such knowledge enables the formation of internal models of the world that we use to plan, make decisions, and act. Recent theorizing posits that mnemonic mechanisms of differentiation and integration – which at one level may seem to be at odds – both contribute to the emergence of structured knowledge. We tested this possibility using fMRI as human participants learned to navigate within local and global virtual environments over the course of three days. Pattern similarity analyses on entorhinal cortex, hippocampus, and ventromedial prefrontal cortex patterns revealed evidence that differentiation and integration work concurrently to build local and global environmental representations, and that variability in integration relates to differences in navigation efficiency. These results offer new insights into the neural machinery and the underlying mechanisms that translate experiences into structured knowledge that allows us to navigate to achieve goals.

## Introduction

Memory is central to who we are, providing the foundation for our sense of self, understanding of the world, and predictions about the future. While we experience life as a series of events, our ability to build knowledge across events – to abstract common themes and infer relationships between events – enables us to construct internal models of the world, or ‘cognitive maps’, that we use to plan, make decisions, and act^1–4^. One goal of memory science is to understand how neural systems integrate information across experiences, building structured knowledge about the world.

Interactions between a network of brain regions in the medial temporal lobe and neocortex are known to support episodic memory, or memory for events. Central to this network is the hippocampus, whose computations allow us to encode and recall specific episodes from the past and to abstract relationships across experiences that share common elements^3,5^. In subserving rapid learning over co-occurring features, the hippocampus plays a critical role in building episodic memories and structured knowledge by binding together inputs from distributed cortical regions, forming conjunctive representations of experiences^3,6,7^. As we learn, neural populations in the hippocampus that represent the traces for individual events can vary in their relations, becoming more distinct (differentiation) or more similar (integration) to each other^8–10^. A central question is how these two seemingly contradictory mechanisms of representational change are employed during the construction of structured knowledge.

One hypothesized hippocampal mechanism is pattern separation, in which the hippocampus creates distinct neural representations for highly similar events^11–13^. Through pattern separation, the hippocampus avoids the blending of similar experiences in memory, reducing across-event interference and forgetting. Tests of the pattern separation hypothesis include functional MRI (fMRI) studies manipulating the extent to which features of events are similar. Such studies have found intriguing evidence for pattern separation when experiences have high overlap or lead to similar outcomes^14–17^, with links to later remembering^18^. Moreover, some fMRI studies have found evidence for pattern differentiation, where event overlap appears to trigger a “repulsion” of hippocampal representations beyond baseline similarity to reduce interference^10,19–24^.

Despite evidence supporting the hippocampus’s role in forming distinct memories for events, decades of work also suggest it is essential for building structured knowledge and forming cognitive maps that capture relations between spatial, perceptual, and conceptual features that were encountered in separate events that share common elements^1,2,4,15,25^. For example, the hippocampus quickly extracts the temporal structure of an environment as defined by the transition probabilities between items in sequences^21,26^, and it represents higher-order structure when there is no variance in transition probabilities between items but an underlying community structure in a sequence^27^. Moreover, it is essential for inferring indirect relationships between items and events^28–30^.

How do overlapping features from separate events lead to integration and generalization? Prior work suggests that retrieval-based learning, recurrent mechanisms within the episodic memory network, and memory replay enable representational integration and allow for the encoding of relationships that have never been directly experienced^27,31–35^. For example, integrative encoding is thought to occur when a new experience (BC) triggers the recall of a related past episode (AB), and the concurrent activation of the remembered and presently experienced episodes results in the formation of an integrated representation (ABC) that can support future inference and generalization^36^. Integrative encoding may relate to other models that rely on recurrent connections between entorhinal cortex (EC) and the hippocampus to recirculate hippocampal output back into the system as new input, allowing for the discovery of higher-order structure at the time of knowledge expression^31–33^. To the extent that the discovered structure is encoded, then across-event relationships are captured in an integrated representation. Moreover, replay within the episodic memory network is also thought to play an important role in updating mnemonic representations^37^. Online replay during periods of awake rest has been shown to integrate multiple events, rather than just replaying a single event^34^. Offline replay during sharp wave-ripples is associated with memory consolidation and updates to neocortical knowledge structures^38–40^.

The building of structured knowledge is not solely the province of the hippocampus. Retrieval-mediated changes in mnemonic representations involve interactions between the hippocampus and ventromedial prefrontal cortex (vmPFC)^25,29,41^. Functional coupling between these regions during learning supports the initial formation of structured knowledge and predicts subsequent memory^42–46^. Over time, vmPFC abstracts and represents commonalities across episodes^25,47,48^, leading some to propose that structured representations are stored in this region^49–52^.

Despite a large and rapidly expanding theoretical and empirical literature, our understanding of how differentiation and integration occur across discrete experiences and contribute to the building of coherent, multi-level structured representations is still far from comprehensive. Additional insights may come from examining these processes during the building of structured spatial knowledge, or maps of the external environment. For example, if one moves to a new city, one often first learns individual routes between navigational goals (i.e., from home to work, home to grocery store). Over time, the episodic memory network links individual routes, building a spatial map that enables planning of novel routes, detours, and shortcuts. Spatial knowledge acquired across learning consists of both local and global representations that are used to different extents based on navigational demands^53^, and the hippocampus and EC are thought to be crucial for the acquisition of such representations. Spatial cells in these regions encode position, head-direction, speed, and other environmental features^54–57^, supporting Tolman’s proposal that navigation is guided by internal cognitive maps and offering insight into how these maps are neurally implemented.

Spatial learning is rich with structure and thus well-suited for exploring the role of differentiation and integration in the formation of local and global knowledge across experiences. Indeed, a previous study in which participants viewed first-person trajectories to real-world destinations found that representations of routes with overlapping sections were differentiated in the hippocampus following learning^20^. Another study observed differentiation of hippocampal patterns for landmarks that were common to multiple virtual cities^52^. By contrast, other studies found evidence of integration for items that share spatial and temporal context^58^ and for objects located in geometrically similar positions across subspaces of segmented environments^59^. More work is needed to characterize experience-driven changes in population-level neural representations of local and global spaces.

Towards this goal, we developed an experimental paradigm that leverages immersive virtual navigation and fMRI to investigate how neural representations of items encountered during goal-directed navigation evolve across learning. Specifically, pattern similarity analyses and linear mixed-effects models characterized experience-driven changes in EC, hippocampus, and vmPFC following “local” and “global” navigation. We hypothesized that we would find evidence for both integration and differentiation emerging at the same time points across learning, as participants build local and global representations of the virtual environment. Moreover, we predicted that early evidence of global map learning during local navigation would depend on integration and predict participants’ ability to subsequently navigate across the environment.

## Results

### Knowledge of local environments

Participants (n=23) completed two behavioral navigation tasks interspersed between three consecutive days of fMRI (Fig.1a). Participants first completed a Local Navigation Task, in which they learned to navigate to five goal locations within each of three distinct local environments –– that is, three oval tracks experienced in a virtual 3D environment from a first-person perspective (Fig.1b). For each track, participants completed four learning runs and six test runs (10 trials/run). At the start of each run, participants were placed on the track and rotated to orient themselves. Each trial began with a fractal cue indicating the navigational goal, followed by the selection of a heading direction and navigation (see Methods; Supplementary Fig.1). We assessed goal-location learning by measuring the proportion of test trials on which participants correctly navigated to the cued location. Navigation was considered accurate if the participant landed within 8 arbitrary units of the goal (36% of the average distance between goals).

**Figure 1.**
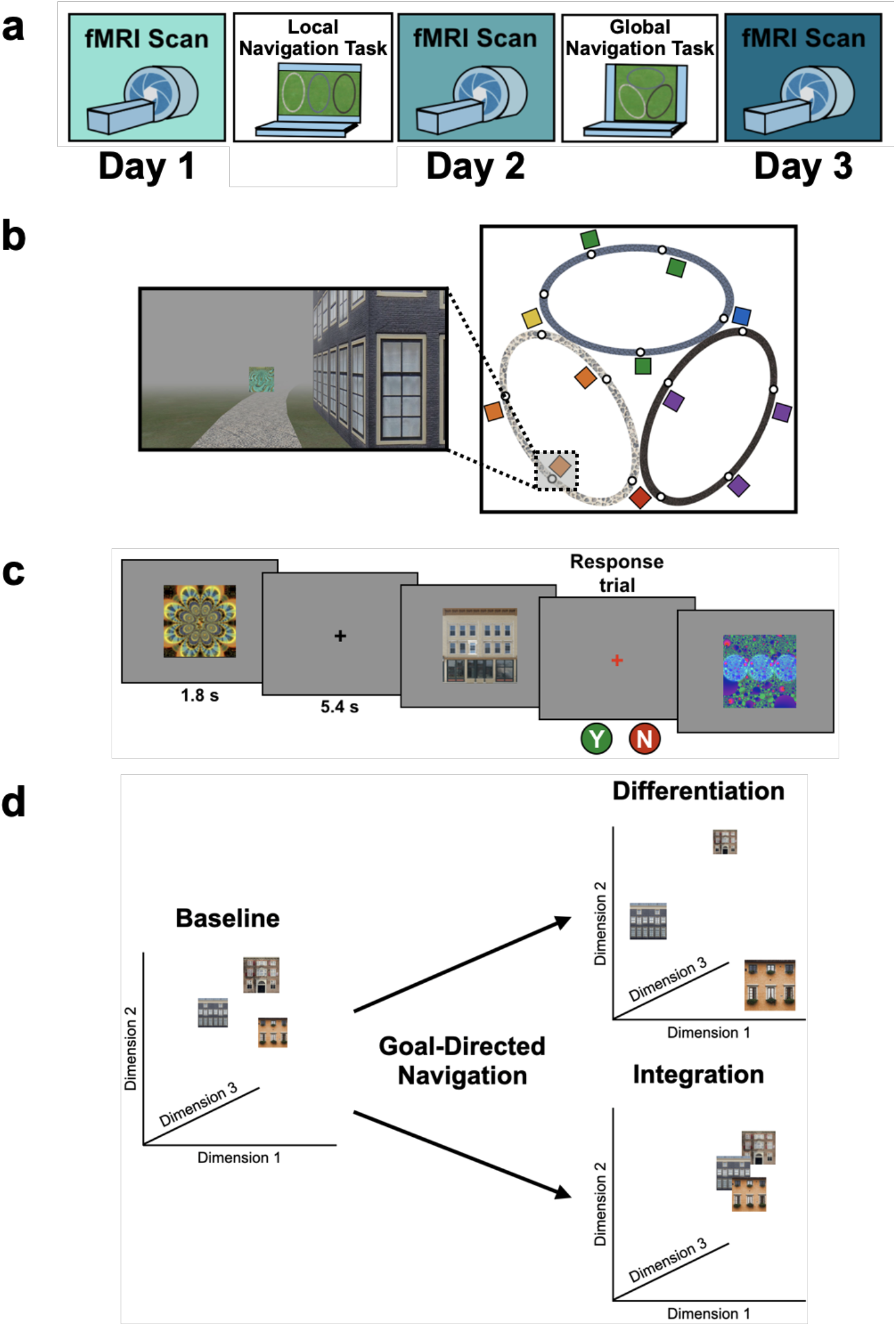
Study design. (A) Overview of the three-day experimental paradigm. (B) First-person view from an example training trial (left); virtual fog limited the distance viewed, ensuring that no two landmark buildings could be seen at any one time. Overhead view of the virtual environment (right). The environment consisted of three oval tracks. Colored boxes indicate the approximate locations of landmarks, circles approximate goal locations for an individual study participant. (C) fMRI paradigm. Participants viewed images of the 12 landmarks and 15 fractals that were used in their unique virtual environment, while performing a perceptual decision-making task. On each trial, the stimulus appeared on a gray background for 1.8s, followed by a fixation cross for 5.4s; participants were instructed to attend to the stimuli and to determine whether a feature of the stimulus was ‘bleached out’; on ‘catch’ trials (8% of trials), a red fixation cross appeared after image offset and participants indicated a response. (D) Schematic illustration of potential representational changes driven by learning. Following goal-directed navigation, the distance between landmarks in neural state-space could have increased (differentiation) or decreased (integration).

Across Day 1 test trials, participants navigated to the correct location on 96% of trials, with performance close to ceiling from the beginning of testing (Fig.2a, top). The proportion of correct trials was significantly above chance (0.25, or 1 of 4 possible goals; mean±SEM: 0.961±0.009; t_22_=82.831, one-sample t-test; p<2.2e^-16^, d=17.27) and did not differ across Day 1 test runs (β=0.003±0.002; t=1.701; p=0.089). To assess whether knowledge of goal locations within local environments was retained overnight, participants completed two additional test runs (Runs 13–14; preceded by two runs of ‘top-up’ learning trials, Runs 11–12). Accuracy on Day 2 (mean±SEM: 0.985±0.004) did not differ from that on the final Day 1 test run (t_22_=1.418; p=0.17, paired t-test).

**Figure 2.**
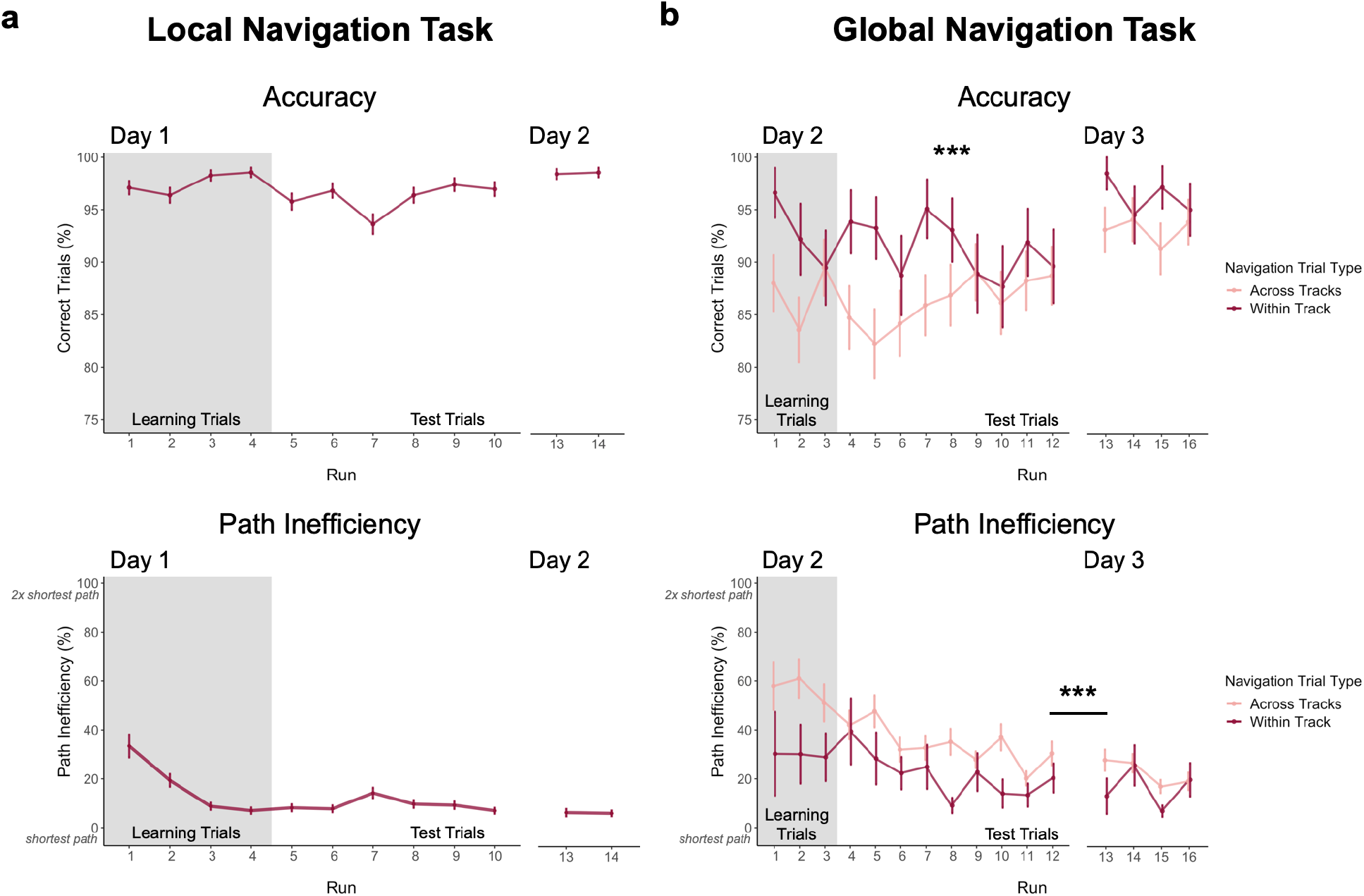
Navigation performance on the Local and Global Navigation Tasks. (A) Local Navigation accuracy on test trials was near ceiling across runs on Day 1 and Day 2 (top). Participants’ navigational efficiency improved over learning trials, and they navigated efficiently across test trials (bottom). (B) During the Global Navigation Task, participants navigated more accurately (top) and efficiently (bottom) on within- vs. across-track trials. Participants were more accurate for within-track trials during learning and during early test runs, but improved on across-track trials over the course of test runs on Day 2 (top). Accuracy improved for both trial types on Day 3, such that performance did not significantly differ between within-track and across-track trials. Participants were significantly more efficient on both trial types on Day 3 relative to Day 2. (*** p < 0.001, paired t-test. Error bars denote SEM. Local Navigation Task, n = 23; Global Navigation Task, n = 21; examined within a single experiment. Source data are provided as a source data file. Learning trials = trials on which the fractal marking the goal location was visible on the track; Test trials = trials on which participants had to rely on memory for the goal location, as the fractals were removed from the track).

Participants were instructed to take the shortest path to a goal. To assess learning of spatial relationships between goals in local environments, we computed a path inefficiency metric that represents a participant’s path length relative to the length of the shortest possible path (see Methods). During Local Navigation (Fig.2a, bottom), path inefficiency improved across the four learning runs (effect of run, β=-8.99±1.262; t=-7.125; p<1.34e^-12^; d=-1.49), was lower on the first test run relative to the first learning run (t_22_=4.272; p=0.0003, paired t-test; d=0.89), and was consistently low across Day 1 test trials (mean±SEM: 9.542±1.773%; no effect of run, β=-0.178±0.378; t=-0.471; p=0.638). As with accuracy, path inefficiency did not differ between Day 2 (mean±SEM: 6.405±1.681%) and the final Day 1 test run (t_22_=-0.619; p=0.542, paired t-test). Together, these data demonstrate that participants quickly acquired precise knowledge of goal locations on the three tracks and the relations between locations within a track, as evidenced by their ability to successfully navigate between goals using nearly the shortest possible path.

### Knowledge of the global environment

Next, 21 of 23 participants completed a Global Navigation Task where the separately learned tracks were connected and participants were required to navigate to goals both within and across tracks. While within-track accuracy on Day 2 was lower in the Global Task (mean±SEM: 0.913±0.031; t_20_=-3.396; p=0.003, paired t-test; d=-0.74, Fig.2b, top) than the preceding Local Task (mean±SEM: 0.985±0.004, Fig.2a, top), performance on the Global Task was above chance for both within-track (mean±SEM: 0.913±0.031) and across-track (mean±SEM: 0.862±0.034) trials (chance=0.07, or 1 of 14 possible goals; within-track trials: t_20_=27.029, one-sample t-test; p<2.2e^-16^, d=5.90; across-track trials: t_20_=23.600, one-sample t-test; p<2.2e^-16^, d=5.15). During Global Navigation, within-track accuracy was higher than across-track accuracy (t_20_=4.073; p=0.0006, paired t-test; d=0.89). However, across-track accuracy showed a trend towards increasing across test runs (effect of run, β=0.007±0.004; t=1.792; p=0.073; no effect of run on within-track trials, β=-0.005±0.004; t=-1.103; p=0.27), such that accuracy did not differ between trial types on the final Day 2 test run (t_20_=-0.029; p=0.977, paired t-test). Before fMRI on Day 3, participants completed four additional test runs of Global Navigation. Accuracy significantly improved overnight for both trial types (within-track mean±SEM: 0.963±0.013, t_20_=2.425, p=0.025, paired t-test, d=0.53; across-track mean±SEM: 0.929±0.025, t_20_=2.841, p=0.01, paired t-test, d=0.62), and did not differ between them on Day 3 (t_20_=1.886, p=0.074, paired t-test; d=0.41).

During Day 2, navigation was less efficient on the Global Task (Fig.2b, bottom) than the preceding Local Task (Fig.2a, bottom). Efficient navigation on across-track trials required a global representation of the virtual environment. During Day 2 learning runs, path inefficiency tended to be greater on across-track (mean±SEM: 59.028±7.124%) than within-track trials (mean±SEM: 31.825±13.392%; t_20_=1.947, paired t-test; p=0.066). Similarly, on Day 2 test runs, navigation was significantly more inefficient on across-track (mean±SEM: 37.86±6.389%) vs. within-track trials (mean±SEM: 23.479±5.764%; t_20_=3.215, paired t-test; p=0.004; d=0.70). However, performance on both trial types improved over the course of Day 2 test runs (within-track trials, effect of run: β=-2.27±1.012; t=-2.244; p=0.025; d=-0.49; across-track trials, effect of run: β=-2.045±0.647; t=-3.163; p=0.002; d=-0.69), such that efficiency did not significantly differ between them during the final Day 2 test run (t_20_=1.480, paired t-test; p=0.155). As with accuracy, efficiency tended to improve overnight for both trial types (Day 3 within-track trials mean±SEM: 17.176±4.222%, t_20_=-3.043, paired t-test, p=0.006, d=-0.66; Day 3 across-track trials mean±SEM: 24.717±4.741%, t_20_=-2.006, paired t-test, p=0.059). Taken together, these data show that participants learned both the local environments (as evidenced by within-track navigation) and the global environment (as evidenced by across-track navigation), achieving high accuracy and efficiency on within- and across-track trials prior to the final scan on Day 3.

### Individual differences in global knowledge

Examination of performance at the participant level revealed striking individual differences in path inefficiency when navigating across tracks in the Global Navigation Task on Day 2 (range=4.219– 107.237%, Q1=17.914%, Q3=54.625%; Fig.5a). Some participants were highly efficient on across-track trials from the beginning, suggesting they had acquired much of the requisite global knowledge during performance of the preceding Local Task. By contrast, others became more efficient at across-track navigation over the course of the Global Task. This variance in when knowledge of the global environment was first evident allowed us to perform an individual differences analysis linking brain activity to behavior that depends on having built global environmental knowledge during Local Navigation (see **Hippocampal representations predict later navigation performance** and Fig.5).

### fMRI assays of learning-driven representational change

Our primary objectives were to characterize representational changes in the human memory network driven by local and global environmental learning. Participants underwent fMRI at three timepoints across the study –– Pre-Learning, Post Local Navigation, and Post Global Navigation (Fig.1a). During fMRI, participants viewed images of the landmark buildings and fractals used in their unique virtual environment, performing a low-level perceptual decision-making task. Of interest was how the neural patterns associated with each landmark and fractal changed as a function of learning (Fig.1d). Accordingly, we extracted voxel-level estimates of neural activity for each stimulus from EC, hippocampus, and vmPFC regions-of-interest (ROIs; see Methods) and used pattern similarity analysis to probe whether learning resulted in representational differentiation or integration as a function of (a) the experienced relations between stimuli in the environment (e.g., same vs. different track; distance within a track) and (b) behavioral differences in when knowledge of the global environment was evident (i.e., evidence of good vs. poor across-track navigation at the outset of the Global Navigation Task). To test neural hypotheses, we fit linear mixed-effects models against pattern similarity, and included fixed effects of interest, nuisance regressors corresponding to the average univariate activation for each stimulus as well as estimates of perceptual similarity, and a standard set of random effects. Complete details of modeling procedures are in supplementary materials.

### Entorhinal cortex learns to separate the three tracks

EC and hippocampus encode a spatial map that may underlie the formation of a cognitive map^2,54–56^. To examine how these regions build structured knowledge, we first asked whether they come to represent the three tracks following learning. We hypothesized that there would be a change in EC and hippocampal pattern similarity for items located on the same track vs. items located on different tracks. To test this hypothesis, we ran models predicting pattern similarity within each region, with scan (Pre-Learning/Day 1, Post Local Navigation/Day 2, and Post Global Navigation/Day 3), stimulus type (landmarks and fractals), and context (same track and different tracks) as predictors. We excluded shared landmarks from these models as they are common to multiple tracks, and restricted pattern similarity comparisons to be within stimulus-type (i.e., landmarks to other landmarks, fractals to other fractals).

An initial model with left and right EC revealed a main effect of hemisphere (*β*=- 0.022±0.004; t=-6.06; p<1.38e^-9^, survived FDR correction; d=-1.43; Supplementary Table 1), and interactions between hemisphere and scan (Day 2>Day 1×hemisphere, β=0.022±0.005; t=4.192; p<2.78e^-5^, survived FDR correction; d=0.99; Day 3>Day 1×hemisphere, β=0.056±0.005; t=10.552; p<2e^-16^, survived FDR correction; d=2.49). Importantly, this model also revealed an interaction between context and scan (Day 2>Day 1×context, *β*=-0.014±0.006; t=-2.05; p=0.041; d=-0.48), such that there was no difference in similarity between within-track and across-track items Pre-Learning (Day 1 same track>Day 1 different track, *β*=0.004±0.004; z-ratio=1.162; p=0.855 adjusted), but within-track items tended to be less similar to each other (i.e., differentiated) following Local Navigation (Day 2 same track>Day 2 different track, *β*=-0.009±0.004; z-ratio=- 2.447; p=0.140 adjusted). No main effect or interactions were observed with stimulus type.

Given the interaction with hemisphere, we ran separate models to examine left and right EC representations of the three tracks as a function of learning. Here, we observed an interaction between context and scan in left EC (Day 2>Day 1×context, *β*=- 0.014±0.006; t=-2.339; p=0.019; d=-0.52; Fig.3a; Supplementary Fig.2a; Supplementary Table 2), such that similarity for same-track vs. across-track items did not differ Pre-Learning, but same-track items were less similar (differentiated) following Local Navigation. Following Global Navigation, pattern similarity remained lower for within-track items, but the interaction between context and scan did not reach statistical significance (Day 3>Day 1×region, *β*=-0.009±0.006; t=-1.457; p=0.145). A similar pattern of findings was observed in right EC, but did not reach statistical significance (Fig.3b; Supplementary Fig.2b; Supplementary Table 3).

**Figure 3.**
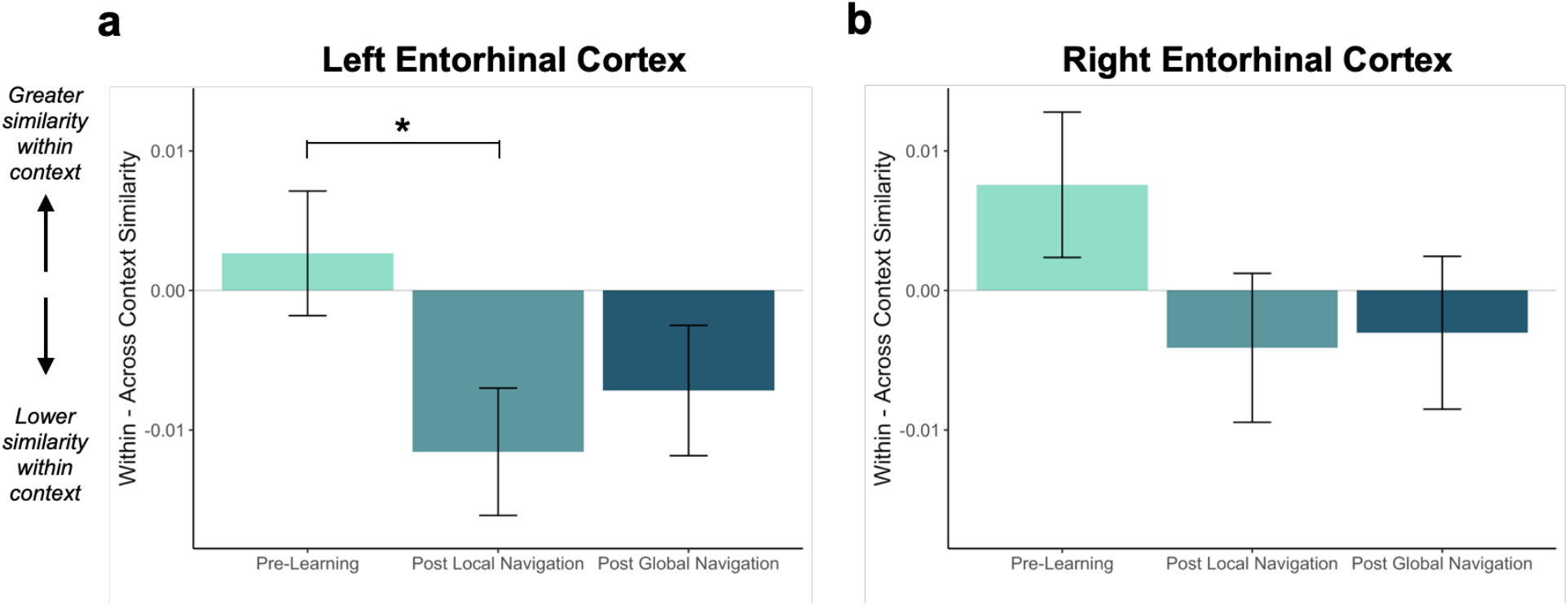
Context representations in entorhinal cortex (EC). Contrast estimates for models in left and right EC. (A) Left EC differentiates items located on the same track following the Local Navigation Task, such that items experienced within the same track became less similar. Following the Global Navigation Task, pattern similarity remained lower for within-track items, but the interaction between context and scan session did not reach statistical significance. (B) A similar pattern of findings was observed in right EC, but interactions between context and scan session did not reach statistical significance. (* p < 0.05. Error bars denote SE of the estimates. Left EC, n = 20; Right EC, n = 18; examined within a single experiment. Source data are provided as a source data file).

Turning to the hippocampus, an initial model that included hemisphere revealed a main effect of hemisphere (*β*=0.006±0.002; t=2.756; p=0.006; d=0.60; Supplementary Table 4), and interactions between hemisphere and scan (Day 2>Day 1×hemisphere, *β*=-0.011±0.003, t=-3.477, p=0.0006, survived FDR correction; d= 0.76; Day 3>Day 1×hemisphere, *β*=0.012±0.003, t=3.795, p=0.0001, survived FDR correction; d=0.83). In contrast to EC, the model also demonstrated a main effect of stimulus type (*β*=0.007±0.003; t=2.422; p=0.015; d=0.53) and an interaction between stimulus type and scan (Day 3>Day 1×stimulus type, *β*=-0.011±0.004; t=-2.604; p=0.009; d=-0.57). The latter interaction reflected a difference in similarity between landmarks and fractals Pre-Learning (Day 1 fractals>Day 1 landmarks, *β*=-0.009±0.002; z-ratio=-4.269; p=0.0003 adjusted) which was no longer present following Global Navigation (Day 3 fractals>Day 3 landmarks, *β*=0.003±0.002; z-ratio=1.248; p=0.813 adjusted).

Given these interactions, we ran models for each hemisphere and stimulus type separately and found evidence in right hippocampus that learning differentiates landmarks located on the same track (Day 3>Day 1×context, *β*=-0.013±0.007; t=-2.014; p=0.044; d=-0.44; Supplementary Fig.3a; Supplementary Tables 5-6). In right hippocampus, Pre-Learning pattern similarity was comparable for within- vs. across-track landmarks. Post Global Navigation, patterns for landmarks on the same track were less similar (differentiated) than those for landmarks on different tracks. No significant interactions between scan and context were observed for fractals (Supplementary Fig.3b; Supplementary Tables 7-8).

To determine whether the effects of learning on representations of the three tracks differed between EC and hippocampus, we included region in a complete model, testing whether pattern similarity in EC for within- vs. across-track items differed from that in the hippocampus. We found a main effect of region (*β*=-0.015±0.002; t=-7.36; p<1.86e^-13^, survived FDR correction; d=-1.74; Supplementary Table 9), as well as interactions between region and scan (Day 2>Day 1×region, *β*=-0.017±0.003; t=-5.757; p<8.59e^-9^, survived FDR correction; d=-1.36; Day 3>Day 1×region, *β*=0.014±0.003; t=4.592; p<4.41e^-6^, survived FDR correction; d=1.08). Importantly, there was no interaction between region, scan, and context (Supplementary Table 9), which suggests that while EC demonstrated significant learning-related within-context differentiation whereas hippocampus did not, statistical support for a functional differentiation between these two regions is absent.

### The hippocampus represents both local and global distance

In studies of episodic memory and spatial navigation, the hippocampus is thought to play a crucial role in disambiguating memories for overlapping events^13,18–20^. Moreover, prior work indicates that spatial distance is reflected in both hippocampal pattern similarity^58^ and univariate BOLD activity^60–62^. We hypothesized that for participants to navigate successfully, they would need to form distinct representations of local environmental landmarks used to aid navigation. We tested whether learning of local environments led to differentiated hippocampal activity patterns by examining pattern similarity for landmarks located on the same track that were nearest neighbors (link distance 1) vs. slightly further away (link distance 2; Fig.4a, left). We ran a model predicting pattern similarity, with scan session and link distance as predictors.

**Figure 4.**
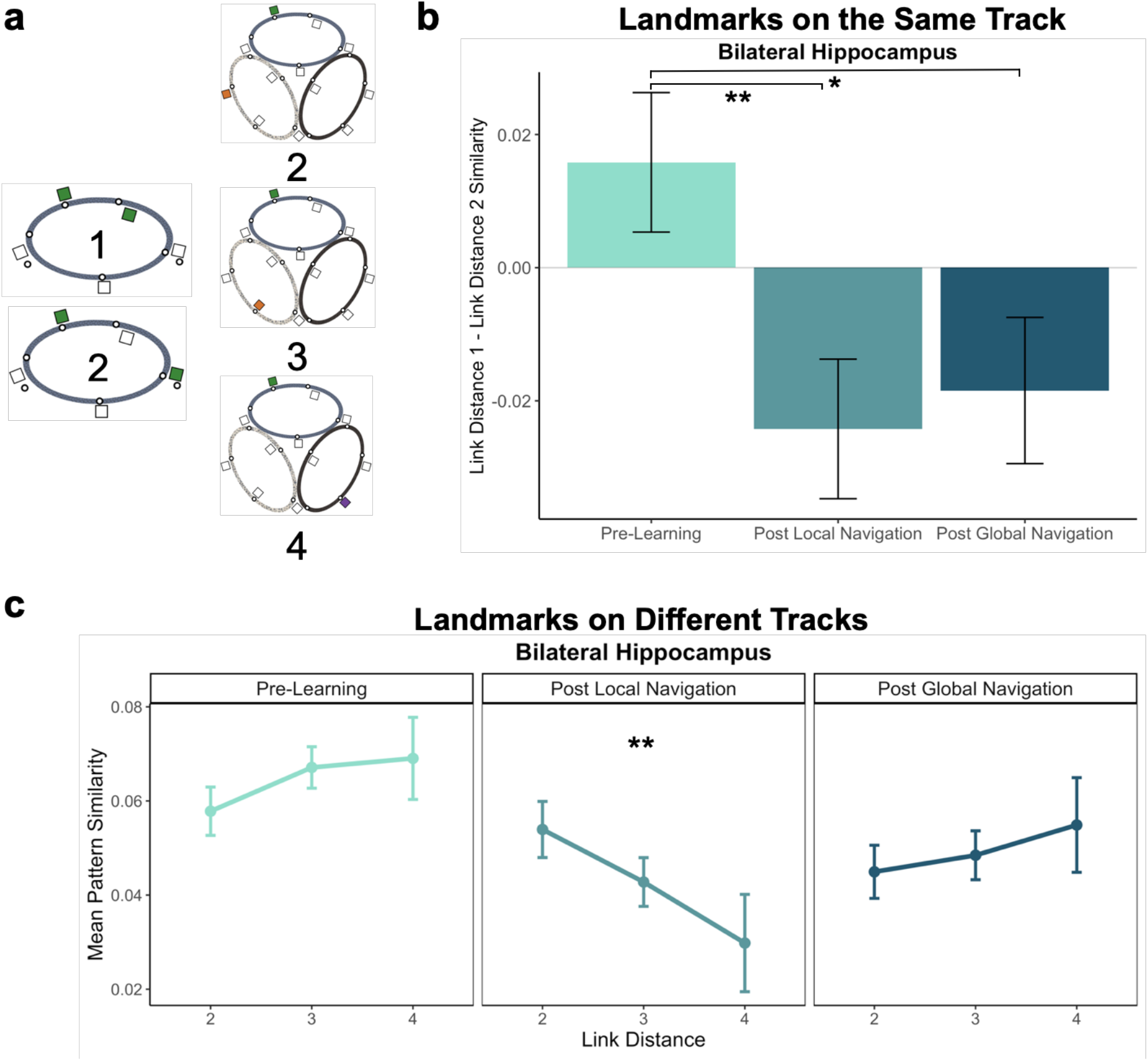
Hippocampal pattern similarity reflects distance in the environment. (A) Examples of landmarks at different link distances on the same track (left) and on different tracks (right). (B) Model contrast estimates show differentiation of hippocampal patterns for landmarks nearest to each other on the same track following the Local and Global Navigation Tasks. (C) Hippocampal pattern similarity for landmarks on different tracks Pre-Learning (left), after the Local Navigation Task (center), and after the Global Navigation Task (right). The interaction between distance and scan session was significant from Pre-Learning to Post Local Navigation, but not to Post Global Navigation. (** p < 0.01, * p < 0.05. Error bars denote SE of the estimates. Day 2 > Day 1, n = 23; Day 3 > Day 1, n = 21; examined within a single experiment. Source data are provided as a source data file).

**Figure 5.**
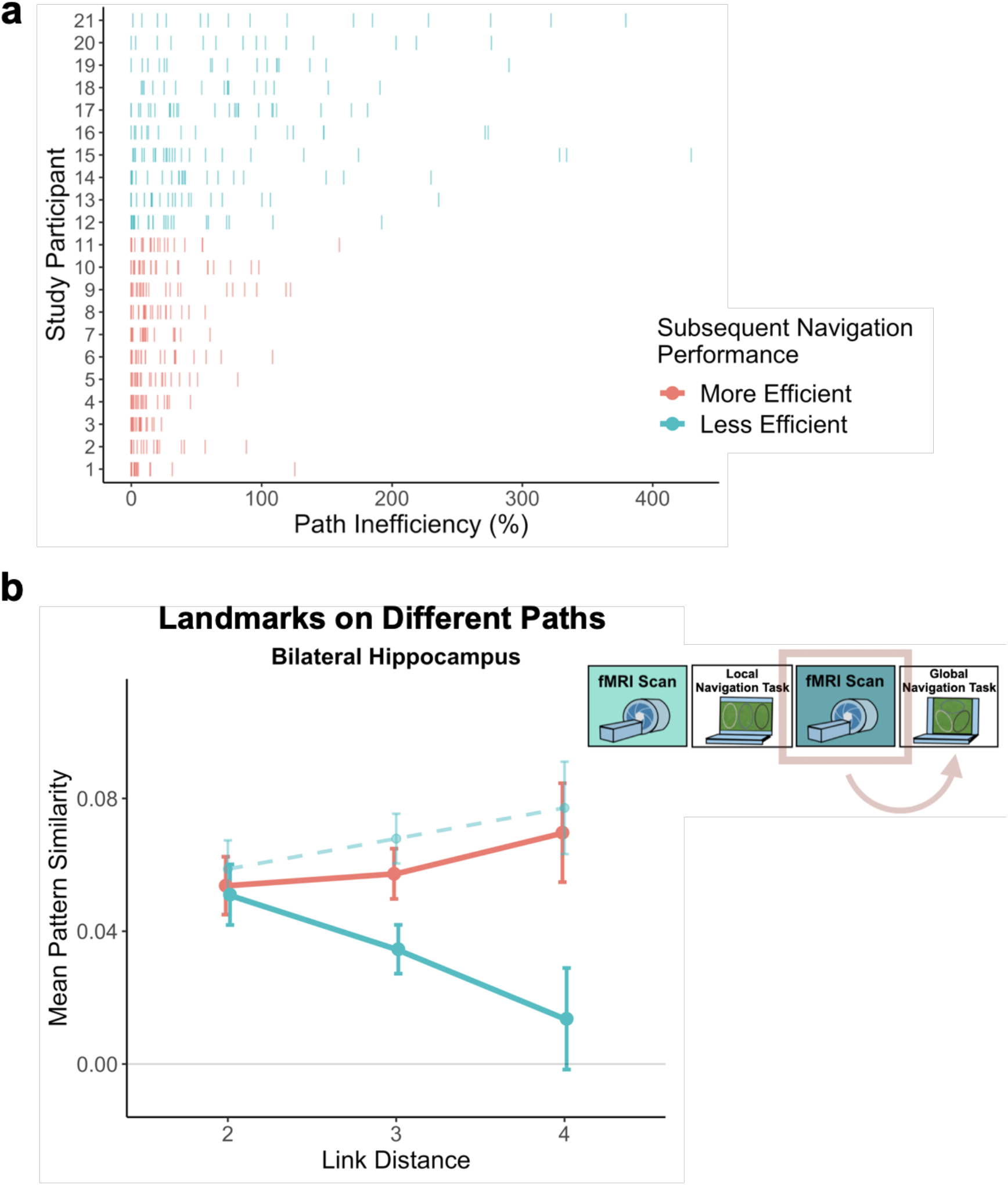
Path inefficiency on the Global Navigation Task varies with across-track distance-related hippocampal pattern similarity after the Local Navigation Task. (A) Path inefficiency (%) for each across-track trial during the first four test runs of the Global Navigation Task, plotted for each participant and colored by performance group. (B) Hippocampal pattern similarity for landmarks on different tracks prior to the Global Navigation Task (solid lines) revealed that the Less Efficient group demonstrated a distance-related effect during Post Local Navigation scanning whereas the More Efficient group demonstrated no effect of distance. Data are split by participants’ subsequent navigation performance as shown in (A); dashed line represents pattern similarity for Less Efficient participants Post Global Navigation (Day 3; see also Supplementary Fig. 6b). (Error bars denote SE of the estimates. More Efficient, n = 11; Less Efficient, n = 10; examined within a single experiment. Source data are provided as a source data file).

An initial model including hemisphere revealed no main effect or interactions with hemisphere (Supplementary Table 10). Importantly, we found interactions between distance and scan (Day 2>Day 1×distance, β=0.04±0.015; t=2.70; p=0.007, survived FDR correction; d=0.56; Day 3>Day 1×distance, β=0.034±0.015; t=2.259; p=0.024; d=0.49; Supplementary Table 11), such that following the Local and Global Tasks, hippocampal patterns were less similar (differentiated) for nearby landmarks vs. landmarks located further apart (Fig.4b).

In addition to encoding local (within-track) spatial representations, navigation may result in the acquisition of a global map of the virtual environment. Although participants performed the Local Task separately on each track, it is possible that the landmarks shared across tracks trigger integration of the tracks, contributing to the encoding of (at least some dimensions of) a global environmental map. We were particularly interested in whether there would be evidence of global knowledge of the environment (i.e., knowledge of relations that span tracks) after the Local Task, prior to any experience navigating across the connected tracks.

To this end, we examined pattern similarity for landmarks located on different tracks at increasing distances, predicting it would scale with distance (Fig.4a, right). To test our prediction, we ran a model predicting pattern similarity between landmarks on different tracks, with scan (Pre-Learning/Day 1, Post Local Navigation/Day 2, and Post Global Navigation/Day 3) and link distance (2, 3, and 4) as predictors. We excluded shared landmarks from this model as they are common to multiple tracks. An initial model that included hemisphere revealed no main effect or interactions with hemisphere (Supplementary Table 12).

Consistent with our prediction, we found an interaction between link distance and scan following the Local Task (Day 2>Day 1×distance, β=-0.02±0.007; t=-2.891; p=0.004, survived FDR correction; d=-0.60), but not the Global Task (Day 3>Day 1×distance, β=-0.003±0.007; t=-0.413; p=0.679; Fig.4c; Supplementary Table 13). In contrast to the distance-related differentiation observed between spatial locations within a track, this interaction reflected that hippocampal pattern similarity for locations across tracks did not vary as a function of distance Pre-Learning, but was higher for closer (vs farther) locations across tracks Post Local Navigation. This finding suggests that some global map knowledge is evident in the hippocampus even though participants had only engaged in within-track navigation. Following Global Navigation, similarity was comparable at all across-track distances, perhaps reflecting acquisition of a more fully integrated global map.

We tested for across-track distance effects within EC as well, given the extensive literature characterizing spatial coding in the region. Contrary to our findings in hippocampus, we found no indication that EC pattern similarity varied as a function of distance, locally or globally (supplementary results; Supplementary Tables 14-18).

### Hippocampal representations predict later navigation performance

The observed negative relationship between hippocampal pattern similarity and the distance between across-track landmarks that emerged after Local Navigation (Fig.4c), but before participants experienced the connected tracks, was quite variable across participants. Another striking feature of the hippocampal global representation is that the relationship with distance observed after Local Navigation was absent following Global Navigation. As speculated above, one possibility is that this latter outcome reflects the presence of more fully integrated global knowledge.

The variability in the observed across-track hippocampal distance effect may reflect that some participants encoded global map knowledge during Local Navigation, whereas others did not (or did so less fully). To the extent that this is the case, this would predict that the distance-related hippocampal pattern similarity effect Post Local Navigation should relate to navigational efficiency at the outset of performing the Global Task. To test this hypothesis, we split participants into two groups – More Efficient (ME) and Less Efficient (LE) – based on the median path inefficiency on across-track trials in the first four test runs of the Global Task on Day 2 (Fig.5a). By definition, the ME group (n=11) was more efficient when navigating across the tracks than the LE group (n=10) on the first four test runs of the Global Task (ME, mean±SEM: 16.892±2.505%; LE, mean±SEM: 72.343±9.835%). Moreover, MEs were more efficient across the preceding learning trials (ME, mean±SEM: 40.124±7.224%; LE, mean±SEM: 79.821±9.018%) and across all test trials (ME, mean±SEM: 16.404±2.474%; LE, mean±SEM: 61.461±8.052%). On Day 3 test trials, MEs continued to outperform LEs (ME, mean±SEM: 12.238±2.541; LE, mean±SEM: 38.443±7.552; t_11_=-3.289, paired t-test, p=0.007; d=-0.99). Importantly, there was a marked improvement in performance for LEs between Days 2 and 3 (t_9_=-5.557, paired t-test, p=0.0004; d=-1.76), while ME performance was not statistically different (t_10_=-1.844, paired t-test, p=0.095), revealing that LEs acquired global map knowledge through performance of the Global Task.

We ran a model predicting pattern similarity Post Local Navigation (Day 2), with group (ME and LE) and link distance (2, 3, and 4) as predictors. Strikingly, we observed an interaction between group and link distance (β=0.025±0.011; t=2.386; p=0.017, survived FDR correction; d=0.52; Supplementary Table 19). Participants who did well from the beginning of the Global Task showed no effect of distance in hippocampal pattern similarity, whereas less efficient navigators showed a negative slope (Fig.5b). As expected, a similar interaction between group and link distance was not observed when the model was fit to data from Day 1 (β=-0.004±0.009; t=-0.46; p=0.646; Supplementary Fig.6a; Supplementary Table 20). Moreover, and consistent with the possibility that a more fully integrated map would result in a null distance-effect, there was no interaction between group and link distance on Day 3 (β=-0.005±0.01; t=-0.472; p>0.637; Supplementary Fig.6b; Supplementary Table 21). Indeed, after extensive experience navigating across the connected tracks, a distance effect was no longer observed across participants (Fig.4c, right) and when restricted to LEs (Fig.5b, dashed).

### Hippocampal-vmPFC interactions

Extant data implicate the vmPFC in learning that requires integration across multiple episodes^25,29,42–47^. Given the learning-related changes observed in the hippocampus, we hypothesized that vmPFC would demonstrate a similar representational similarity structure. To test this, we first asked whether vmPFC pattern similarity for landmarks at different distances significantly differed from the hippocampus by running a complete model that included data from both regions. Here, we found no effect of region or interactions between region, scan, and distance (Supplementary Table 22), indicating that the similarity structure was not significantly different between regions. By contrast, pattern similarity in both the hippocampus and vmPFC differed from that of a visual control region (see supplemental results; Supplementary Table 23). However, in contrast to hippocampus, models predicting pattern similarity for landmarks at different link distances in vmPFC revealed no interactions between link distance and scan for within-track (Supplementary Fig.7a; Supplementary Table 24) or across-track landmarks (Supplementary Fig.7b; Supplementary Table 25).

While distance-related effects on vmPFC and hippocampal pattern similarity did not significantly differ, such effects were significant in hippocampus but not in vmPFC. To further test whether distance-related similarity structures were similar between the regions, we correlated the similarity matrices for vmPFC and the hippocampus at the individual subject level, and compared this correlation to one with a visual region acting as a control. Here, we found that the vmPFC-hippocampus correlation was significantly greater than the vmPFC-visual cortex correlation in all scan sessions (Pre-Learning: t_22_=-5.979; p<5.2e^-6^, paired t-test; d=-1.25; Post Local Navigation: t_22_=-3.007; p=0.006, paired t-test; d=-0.63; Post Global Navigation: t_20_=-5.499; p<2.205e^-5^, d=-1.20; Fig.6). However, there was no interaction between region and scan (Supplementary Table 25).

**Figure 6.**
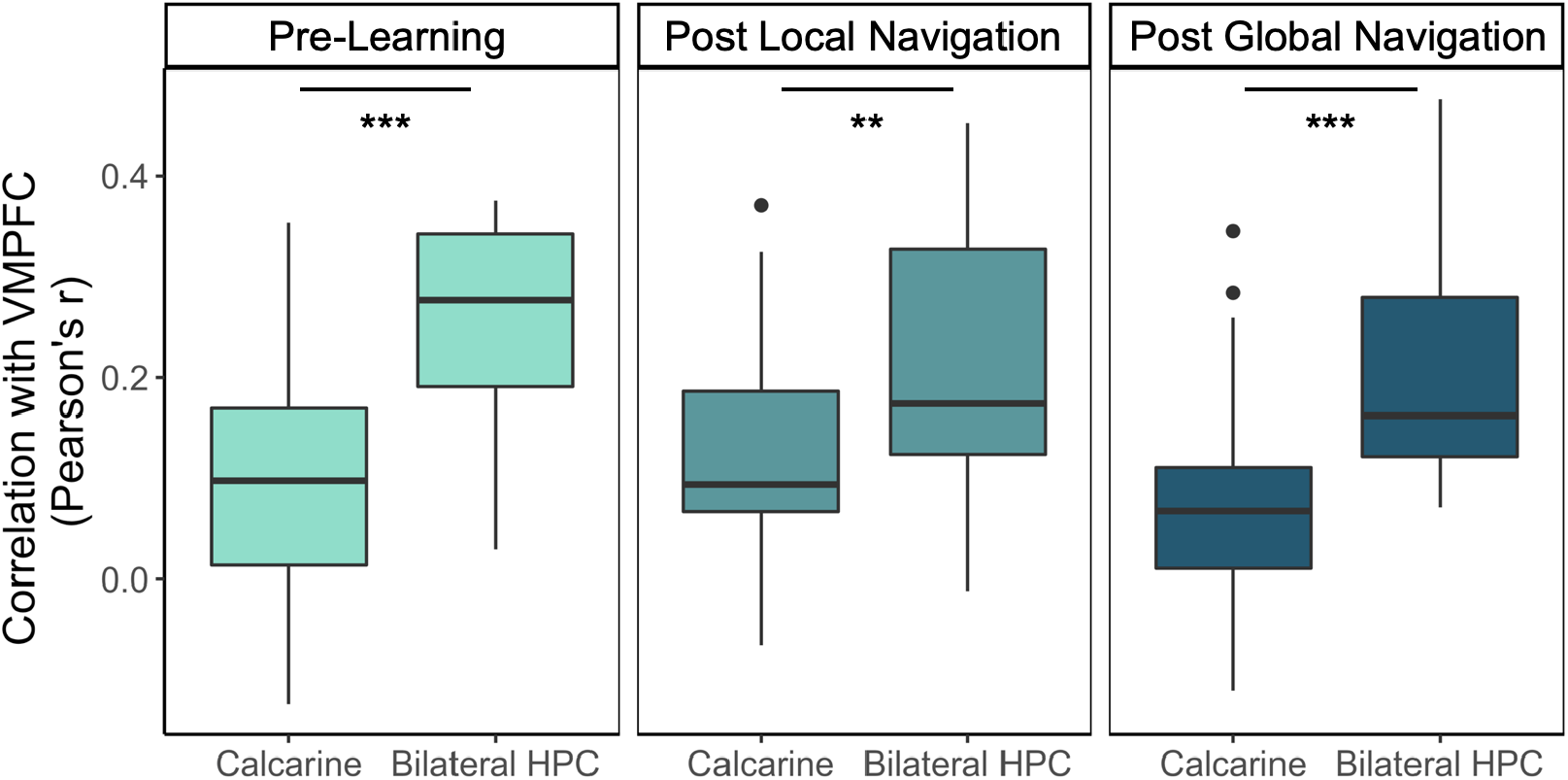
The neural similarity structure for landmark buildings at different distances is more similar between vmPFC and hippocampus (bilateral HPC) than vmPFC and a visual control region (calcarine). Boxplots display median values and data quartiles, whiskers extend to 1.5 times the minima/maxima in the absence of outliers. (*** p < 0.001 ** p < 0.01, paired t-test. Pre-Learning, Post Local Navigation, n = 23; Post Global Navigation, n = 21; examined within a single experiment. Source data are provided as a source data file).

We suspected that the lack of learning-related effects in vmPFC may have been due to our use of an overly inclusive ROI mask, so we generated an 8mm-radius spherical ROI around the peak voxel reported in an fMRI study of declarative memory consolidation^50^. We again found no significant distance effects using this smaller ROI, but did observe a trend for a context effect consistent with our findings in EC and hippocampus (Supplementary Fig.8, Supplementary Tables 27-29).

## Discussion

The episodic memory network builds structured knowledge across experiences to form internal models of the world that enable planning, decision-making, and goal-directed behavior. In this study, we investigated how humans build structured knowledge in immersive, goal-directed tasks. Results revealed that (a) learning restructures representations in the hippocampus and EC, reflecting the structure of the virtual environment; (b) the hippocampus begins to build a representational structure extending beyond directly experienced transitions; and (c) changes in the similarity structure of hippocampal representations relate to subsequent navigation performance.

We first characterized learning on two behavioral navigation tasks. In the Local Navigation Task, participants quickly learned goal locations along individual tracks and navigated between goals using the most efficient paths (Fig.2). Prior to the Global Navigation Task, participants were unaware that the tracks would be connected and that they would be required to navigate across tracks to reach cued goals. Nonetheless, some were able to navigate efficiently from the outset of Global Navigation, suggesting they had already formed a global representation of the environment; by contrast, others required extensive experience to do so (Fig.5a). Our findings are consistent with behavioral research examining individual differences in route integration and cognitive map formation. Prior work characterized navigators into groups of “integrators”, “nonintegrators”, and “imprecise navigators”, suggesting that while some individuals build accurate internal maps, others may rely on fragmented maps or route-based strategies during navigation^63–65^. Importantly, individual differences in navigation inefficiency enabled us to probe variability in neural representations and relate neural pattern similarity to behavior. Our work provides an important extension of prior studies in that it identifies potential neural substrates of individual differences in cognitive map formation.

At the neural level, we investigated whether mnemonic regions come to represent the three local tracks following navigation by comparing pattern similarity for items located on the same track vs. different tracks. Motivated by recent studies finding lower hippocampal pattern similarity for events with overlapping vs. distinct features^15,18– 20,24^, we expected to observe changes in hippocampal pattern similarity for within-track items following local learning. Indeed, we found evidence for differentiation in EC: items on the same track elicited less similar activity patterns when compared to patterns for items on different tracks following local learning (Fig.3). While we observed a quantitatively similar pattern of effects for landmarks in the hippocampus, they were not statistically significant (Supplementary Fig.3a).

Several factors may explain why we did not find stronger hippocampal differentiation effects. The demands of our navigation tasks differ substantially from those of prior tasks eliciting strong differentiation effects^15,16,19,20,66^. In prior studies, participants explicitly learned overlapping associations that had to be distinguished at a later test; for instance, the same item was paired with two different associates, or similar items or routes led to distinct outcomes. In the present study, overlapping features were incidental to task demand; thus individuals may have adopted different navigational strategies with varying consequences for within-track item pattern similarity (for instance, some participants may have learned the sequence of fractals for each track, while others may have formed associations between fractals and nearby landmarks). Tracks had a number of features in common, including shared visual features, a similar spatial layout, and common landmarks. Further, our findings from across-track analyses examining individual differences in navigation suggest that some participants began to build an integrated, global representation of the environment prior to the Global Task, with associated effects in the hippocampus. Given these dynamics, additional variance inherent in our paradigm may explain the nature of the observed effects in hippocampus and EC.

While we did not observe particularly strong differentiation of hippocampal patterns for all items located on the same track, we hypothesized that participants would need to form distinct representations of local environmental landmarks to perform well. Accordingly, we examined local spatial representations by comparing the similarity between nearest neighboring landmarks to second nearest neighbors on the same track, and found that hippocampal patterns for nearest neighboring landmarks were differentiated following both Local and Global Navigation (Fig.4b). This finding is consistent with the idea that mnemonic representations – and in particular, representations of large-scale spaces – are hierarchical^67–69^. That is, we found clearly differentiated representations of local space, whereas representations of track context or of the spatial relationships between the tracks were perhaps more coarse.

We next investigated global representations of map knowledge within the virtual environment. Here, we examined hippocampal pattern similarity for landmarks encountered on different tracks at increasing distances in the virtual environment, expecting to see the emergence of a relationship between hippocampal pattern similarity and distance. We were particularly interested in whether this relationship would be observed after the Local Task, which would suggest that participants were starting to build a global map of the environment prior to direct experience navigating the connected tracks.

Our results provide support for this prediction – we found a significant interaction between distance and scan (Pre-Learning vs. Post Local Navigation; Fig.4c, center). Notably, we also observed significant variability in the slope of this function across individuals. We hypothesized that this variability in the neural data might relate to the behavioral variability we observed on the subsequent Global Navigation Task, so we split participants into two groups based on their path inefficiency on the earliest Global Navigation trials and found a significant interaction between group and distance (Fig.5b). The interaction between distance and scan (the negative slope in the center panel of Fig.4c) was driven by participants who were less efficient at the outset of Global Navigation.

Our results suggest that a negative distance-related similarity function reflects restricted learning of the global map, hindering performance on the Global Task. While our findings cannot definitively speak to what this negative relationship represents, we believe there are two possibilities. First, LE participants may have differentiated more distal landmarks, with such differentiation being less effective in supporting across-track navigation. While a potential interpretation, our results are more compatible with an alternative interpretation centered on mnemonic integration and the building of a more global cognitive map. Specifically, the data suggest that, prior to Global Navigation, LEs had integrated only the nearest landmarks located on different tracks (link distance 2), while MEs had integrated all across-track link distances. Accordingly, we observed an increase in pattern similarity for across-track landmarks at link distances 3 and 4 in LEs only after they had acquired substantial global map knowledge through Global Navigation (Fig.5b, dashed line, right; Supplementary Fig.6b).

Replay and recurrence within the hippocampal network is thought to enable the hippocampus to learn relationships between items that have never been directly experienced together^29,31,32,34,35^. The present study provides complementary evidence that the hippocampus begins to build a cognitive map extending beyond directly experienced transitions in a goal-directed task. Overlapping features within the virtual environment (the landmarks shared between tracks) provide the hippocampus with the links it needs to begin building a global map, despite participants not yet experiencing the connected environment. Crucially, not all participants demonstrated integration early in the Global Task, suggesting that cognitive map formation may not be a ‘default’ that incidentally emerges during performance of immersive, goal-directed local navigation. An open question is: How does this variability in whether and how individuals build global knowledge of the environment during local navigation relate to individual differences in spatial and mnemonic processing, in navigational strategy, and/or in other cognitive and contextual factors^64,65,70^? The current study was limited in its investigation of individual differences due to sample size. Future work is needed to characterize the differences between performance groups and examine how representations of individual tracks and particular spatial positions within tracks covary with performance on subsequent global navigation. Moreover, while some of our findings do not survive correction for multiple comparisons, and thus should be interpreted with caution, the key integration and differentiation findings within the hippocampus remain significant following correction.

The present findings appear to diverge from work finding hippocampal pattern similarity scaled with perceived spatiotemporal distance in a virtual environment, such that patterns for nearby items were integrated^58^. In that study, participants were scanned before and after a task with a duration of ∼80min; thus, the findings may reflect hippocampal representations at an earlier stage of learning. In the present paradigm, fMRI took place 21.5–31hrs after extensive navigational practice, raising the possibility that hippocampal differentiation requires extensive learning, potentially including learning that takes place via replay and consolidation processes occurring during sleep. Future studies are necessary to pinpoint when integration occurs across the broader global map.

Mnemonic integration depends in part on retrieval and interactions between the hippocampus and vmPFC^41,46,48,52^. We were interested in whether vmPFC pattern similarity would mirror our findings in hippocampus, and/or how vmPFC would come to represent the virtual environment, given its proposed role as a storage site for integrated memory representations^49–51^. Potentially consistent with what we observed in hippocampus and EC, we observed trend-level evidence that vmPFC comes to represent the three tracks after local learning. Yet, while we did not find statistically significant evidence that vmPFC comes to represent distance in the global environment, pattern similarity did not differ significantly from what we observed in hippocampus. Correlations between similarity matrices in vmPFC and the hippocampus were significantly greater than those between vmPFC and a visual control region; but as this was true at all three scans, these relationships do not reflect learning-related representational changes. It remains possible that vmPFC may represent features of the environment that were not explicit to our modeling approach.

An emergent body of work is advancing understanding of fundamental mechanisms in the medial temporal lobe and connected cortical structures that build knowledge across events. Part of this work documents pattern separation and pattern differentiation in the hippocampus, which serves to minimize confusion between related memories. At the same time, other theoretical and empirical findings are elucidating the role of these structures in mnemonic integration and cross-event generalization. It has been unclear whether and how these two seemingly opposing mechanisms –– differentiation and integration –– work together to facilitate the emergence of structured knowledge. Here, we provide complementary evidence that both differentiation and integration occur concurrently within the hippocampus and can emerge at the same points in learning. Our data suggest that hippocampal differentiation provides a structure necessary for participants to distinguish between nearby locations, while integration processes serve to build a global map of the environment. Moreover, integration relates to individual differences in the ability to efficiently navigate between locations in the global map that have not been previously traversed. Future research promises to reveal whether individual differences in global knowledge building are strategic, relate to differences in other neural systems, and/or are differentially sensitive to dysfunction or disease.

## Methods

### Participants

Thirty-three participants gave informed written consent, in accordance with procedures approved by the Stanford University Institutional Review Board. Nine dropped out before completing the three-day study (two felt motion sick, two experienced unrelated illness, and five did not want to continue past Day 1), and data from one participant were excluded for failing to respond during the fMRI task. The final sample consisted of 23 right-handed participants (mean age=22.91yrs, SD=5.03, range 18–35; 14 females) with normal or corrected-to-normal vision and no self-reported history of psychiatric or neurological disorders. Day 3 data were excluded for two of these participants (one due to scanner-related issues that prevented data acquisition, and one who reported falling asleep during scanning). Thus, Global Navigation Task/Day 3 analyses were conducted with 21 participants. Participants received compensation for their participation ($20/hr).

### Paradigm overview

Participants completed an experimental paradigm where they learned to navigate to goal locations within a virtual environment. The three-day study included two behavioral navigation tasks –– Local Navigation Task and Global Navigation Task –– and three fMRI scan sessions (Fig.1a) that assayed changes in hippocampal and neocortical representations across learning. The virtual environment contained three oval-shaped tracks of distinct texture and color (Fig.1b). Individual tracks contained five goals, initially marked by unique fractal images. Tracks also contained unique landmark buildings that participants could use to guide their navigation, as there were no distal cues in the virtual environment. Landmarks were located near (but not at) goals, with the distance between them varying for each location and participant. Importantly, each track had one landmark in common with each of the other two tracks.

Participants first completed the Local Navigation Task, where they learned to navigate to goals on each of the three tracks separately, with fog serving to hide others from view (at any given point on a track, only one landmark and goal was visible in the field of view). The next day, participants completed the Global Navigation Task, where the separately learned tracks were connected and participants were required to navigate to goals both within and across tracks (Fig.1b, right). Participants were not informed and not aware that the tracks would be connected prior to the start of the Global Task, despite the shared landmarks. Efficient navigation on the Global Task required knowledge of the spatial relationships between locations in the global environment both within and across tracks. For complete details of task stimuli, see supplementary methods.

### Virtual Navigation Tasks

The navigation tasks were developed with Panda3D (1.9.4), a python-based open-source gaming engine, and the PandaEPL library^71^. For all navigation tasks, participants were instructed to navigate to goals using the shortest possible path. Each behavioral run contained 10 navigation trials. At the start of each run, participants were placed at a location on the path and slowly rotated 360 degrees to orient themselves (6s). Trials then proceeded as follows: 1) a fractal cue appeared onscreen (1s) indicating the goal to which the participant should navigate; 2) participants chose their heading direction and 3) navigated to the cued goal, pressing the spacebar at arrival; 4) feedback appeared onscreen revealing whether the participant was at the correct location and had navigated via the shortest path (2s); and 5) the camera panned down and a fixation cross appeared (1s) before the next trial began (Supplementary Fig.1). Participants navigated in the environment by pressing the forward arrow key and adjusted their heading direction using the left and right arrow keys. Movement was fixed along the centerline of the paths and movement speed was held constant. Traveling around one individual path required approximately 30s.

### Local Navigation Task

During the Local Navigation Task, participants learned to navigate to goals on the individual tracks separately, with the other tracks hidden from view. After fMRI scanning on Day 1 (see fMRI task below), the Local Task began with four runs of learning trials on track 1. On learning trials, fractals were visible at goals to allow for learning of their locations within the track environment. The participants then completed two runs of test trials, where fractals were no longer visible and participants had to rely on memory to navigate successfully. This procedure was repeated for track 2 and track 3. After completing two runs of test trials on each track, participants began additional interleaved blocks of test trials, switching between tracks until six test runs (60 trials) were completed for each of the individual tracks. In total, participants navigated from every goal on a track to every other goal on that track, several times. Track order was randomized and counterbalanced across participants. After Day 1, participants returned 24hrs (range=21–31.5hrs) later for Day 2, which started with two additional runs of Local Navigation Task test trials on each track prior to fMRI on Day 2. These two runs provided an assay of whether knowledge of the individual tracks had been retained overnight.

### Global Navigation Task

After fMRI on Day 2, participants began the Global Navigation Task. Here, the three tracks were connected and participants were required to navigate to goals both within and across the different tracks. In this task, common landmarks served as linking points within the larger environment (Fig.1b). Participants first completed three runs of learning trials, where fractals were again visible at goals. While the goal locations did not change between the Local and Global Navigation Tasks, learning trials allowed participants to become accustomed to moving across tracks and gave them an opportunity to orient themselves to the larger environment. Participants then completed nine runs of test trials, where fractals were no longer visible in the environment. For 30 of 90 test trials, participants were cued to navigate to a goal along the track on which they were already located. For the other 60, participants were required to navigate to a different track. After Day 2, participants returned 24hrs (range=15.5–28.5hrs) later for Day 3, which started with four additional runs of Global Navigation test trials prior to fMRI on Day 3. These four runs provided an assay of whether spatial knowledge had been retained overnight.

### Behavioral data analysis

To assay whether participants successfully learned goal locations, we computed percent correct as the ratio of the number of trials where participants ended navigation within 8 a.u. of the virtual goal location (defined as a correct response) over the number of trials attempted. To assay whether participants had learned spatial relationships between the locations in the environment, we computed a path inefficiency metric by dividing path length by the length of the shortest possible path to the goal, subtracting 1, and multiplying by 100 to express as a percent. Thus, a path inefficiency of 0% would indicate that the participant took the shortest path possible, and a path inefficiency of 100% would indicate that their path was twice the length of the shortest possible path to the goal.

### fMRI task

Participants underwent fMRI scanning prior to any learning (Day 1), after the Local Navigation Task (Day 2), and after the Global Navigation Task (Day 3). During each fMRI session, participants viewed images of the 12 landmarks and 15 fractals that were used in their unique virtual environment. Stimuli were presented 12 times per scan session (3 repetitions/run x 4 runs). Within each scan, the stimuli appeared in mini-blocks such that participants saw all 27 images prior to seeing any repeated, with third repeats appearing only after all stimuli had appeared twice. Images were pseudo-randomized within each mini-block such that the same image could not appear within three steps of itself at the block transitions. Moreover, no first order stimulus-to-stimulus transitions were repeated within a scan session.

On each trial, the stimulus appeared on a gray background for 1.8s, followed by a fixation cross of 5.4s (intertrial interval; Fig.1c). To ensure that participants paid close attention to the images, they performed a visual anomaly detection task in which they were asked to report whether a feature of an image was ‘bleached out’ on trials when a red fixation cross appeared after image offset^72^. Only a small proportion of stimulus presentations were response trials (∼8%). The probability of a response trial having a bleached feature was 0.5. Response trials were excluded from all further analyses.

### MR data acquisition

Whole-brain imaging data were acquired on a 3T GE Discovery MR750 MRI scanner (GE Healthcare) using a 32-channel radiofrequency receive-only head coil (Nova Medical). Functional data were acquired using a 3-band echo planar imaging (EPI) sequence (acceleration factor=2) consisting of 63 oblique axial slices parallel to the long axis of the hippocampus (TR=1.8s, TE=30 ms, flip angle=75°, FOV=220mm×220mm, voxel size=2×2×2mm^3^). To correct for distortions of the B0 field that may occur with EPI, two B0 field maps were acquired before every functional run, one in each phase encoding direction, with the same slice prescription as the functional runs. Structural images were acquired using a T1-weighted (T1w) spoiled gradient recalled echo structural sequence (186 sagittal slices, TR=7.26ms, FoV=230mm×230mm, voxel size=0.9×0.9×0.9mm^3^). The MR data collection techniques closely mirrored procedures in^23^. MR data were preprocessed according to standard pipelines (see supplementary methods).

### fMRI analysis

Prior to fMRI analyses, we removed the first 6 TRs in each run. For each scan session, we built general linear models (GLMs) for even and odd runs, which were regressed against preprocessed fMRI data at the voxel level. To obtain stimulus-level beta estimates for brain activity, each stimulus (i.e., landmark or fractal) was represented by a single regressor, time-locked to when it appeared onscreen in odd or in even runs. Each event was modeled as an epoch lasting for the duration of stimulus presentation (1.8s) and convolved with a canonical hemodynamic response function. A separate regressor was included for response trials to exclude them from further analysis. Response trials were modeled as epochs lasting for the duration of the stimulus presentation and subsequent response period (7.2s). GLMs also included nuisance regressors marking outlier TRs (DVARS>5 or FD>0.9 mm from previous TR), a run regressor, and regressors generated by fMRIPREP representing TR-level 6-dimensional head movement estimates, framewise displacement (mm), and low frequency components for temporal high-pass filtering. GLMs yielded estimates of voxel-level brain activity for each stimulus during odd and even runs of each scan session. Modeling was performed with SPM12 (Wellcome Trust Centre for Neuroimaging) and custom Matlab (vR2017b) routines.

Primary analyses were performed using an *a priori* region-of-interest (ROI) approach targeting the hippocampus, EC, and vmPFC. *ROI segmentation*. Bilateral hippocampal, entorhinal, parahippocampal, and perirhinal cortical ROIs were manually delineated on each participant’s high-resolution T1-weighted structural image using established procedures^73^. Due to low TSNR (<28), selected scans from 3 participants were excluded from analyses in right EC, and 1 scan from 1 participant was excluded from analyses in left EC. We obtained a vmPFC mask from a Neurosynth parcellation^74^, and transformed the mask into native space for each participant. We also created a smaller 8mm spherical vmPFC mask using the peak voxel reported in Takashima et al. (2006)^50^. ROIs were resampled, masked to exclude voxels outside the brain, and aligned with functional volumes. Finally, we used a visual ROI defined by FreeSurfer’s automated segmentation procedure as a control for our analyses. The mask for this ROI was defined as a conjunction of FreeSurfer’s pericalcarine and calcarine sulcus regions in both hemispheres.

The activity pattern for each stimulus was quantified as a vector of multi-voxel normalized betas by dividing the original betas by the square root of the covariance matrix of the error terms from the GLM estimation^75^. Separately for each participant, ROI, and scan session, we computed pattern similarity by correlating activity patterns between even and odd runs. Pattern similarity analyses were conducted in Python (v2.7).

### Statistical analyses

Statistical analyses were implemented in R (v3.6.3). For neural analyses described above, we implemented linear mixed-effects models using the lme4 and lmerTest statistical packages^76,77^. Correlations, our dependent variable, were Fisher transformed to follow a normal distribution. Models included fixed effects of interest (i.e., scan session and link distance), fixed effects corresponding to the average univariate activation for each of the stimuli, fixed effects estimating perceptual similarity in V1 and IT cortex, and a standard set of random effects, including a random intercept modeling the mean subject-specific outcome value, as well as random slope terms modeling subject-specific effects of independent variables of interest (i.e., scan session and average univariate activation). We included average univariate activation as a regressor to limit the possibility that overall activation differences within an ROI would impact pattern similarity^78^. Comprehensive modeling details can be found in the supplementary methods.

FDR (false discovery rate) correction controlled for multiple comparisons. Unless otherwise specified, all P-values reported were uncorrected. If applicable, we note whether the reported results survived FDR correction. Cohen’s *d* effect sizes were computed for t-values as 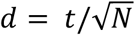.

## Acknowledgements

This project was supported by The Marcus and Amalia Wallenberg Foundation and The Stanford Center for Cognitive and Neurobiological Imaging.

## Data availability

Raw MRI data and behavioral data will be available on OpenNeuro. Analytical code and data for analyses are available at https://github.com/coreyfernandez/RID.

## References

1. Tolman, E.C. Cognitive maps in rats and men. Psychol. Rev. 55, 189–208 (1948).

2. O’Keefe, J. & Nadel, L. The hippocampus as a cognitive map. (Clarendon Press ; Oxford University Press, 1978).

3. McClelland, J. L., McNaughton, B. L. & O’Reilly, R. C. Why there are complementary learning systems in the hippocampus and neocortex: insights from the successes and failures of connectionist models of learning and memory. Psychol. Rev. 102, 419–457 (1995).

4. Epstein, R. A., Patai, E. Z., Julian, J. B. & Spiers, H. J. The cognitive map in humans: spatial navigation and beyond. Nat. Neurosci. 20, 1504–1513 (2017).

5. Eichenbaum, H. Hippocampus: cognitive processes and neural representations that underlie declarative memory. Neuron 44, 109–120 (2004).

6. Teyler, T. J. & DiScenna, P. The hippocampal memory indexing theory. Behav. Neurosci. 100, 147–154 (1986).

7. Libby, L. A., Hannula, D. E. & Ranganath, C. Medial temporal lobe coding of item and spatial information during relational binding in working memory. J. Neurosci. Off. J. Soc. Neurosci. 34, 14233–14242 (2014).

8. Ritvo, V. J. H., Turk-Browne, N. B. & Norman, K. A. Nonmonotonic Plasticity: How Memory Retrieval Drives Learning. Trends Cogn. Sci. 23, 726–742 (2019).

9. Brunec, I. K., Robin, J., Olsen, R. K., Moscovitch, M. & Barense, M. D. Integration and differentiation of hippocampal memory traces. Neurosci. Biobehav. Rev. 118, 196–208 (2020).

10. Wammes, J., Norman, K. A. & Turk-Browne, N. Increasing stimulus similarity drives nonmonotonic representational change in hippocampus. eLife 11, e68344 (2022).

11. O’Reilly, R. C. & McClelland, J. L. Hippocampal conjunctive encoding, storage, and recall: Avoiding a trade-off. Hippocampus 4, 661–682 (1994).

12. Leutgeb, J. K., Leutgeb, S., Moser, M.-B. & Moser, E. I. Pattern separation in the dentate gyrus and CA3 of the hippocampus. Science 315, 961–966 (2007).

13. Yassa, M. A. & Stark, C. E. L. Pattern separation in the hippocampus. Trends Neurosci. 34, 515–525 (2011).

14. Bakker, A., Kirwan, C. B., Miller, M. & Stark, C. E. L. Pattern separation in the human hippocampal CA3 and dentate gyrus. Science 319, 1640–1642 (2008).

15. Schlichting, M. L., Mumford, J. A. & Preston, A. R. Learning-related representational changes reveal dissociable integration and separation signatures in the hippocampus and prefrontal cortex. Nat. Commun. 6, 8151 (2015).

16. Kyle, C. T., Stokes, J. D., Lieberman, J. S., Hassan, A. S. & Ekstrom, A. D. Successful retrieval of competing spatial environments in humans involves hippocampal pattern separation mechanisms. eLife 4, e10499 (2015).

17. Ballard, I. C., Wagner, A. D. & McClure, S. M. Hippocampal pattern separation supports reinforcement learning. Nat. Commun. 10, 1073 (2019).

18. LaRocque, K. F. et al. Global Similarity and Pattern Separation in the Human Medial Temporal Lobe Predict Subsequent Memory. J. Neurosci. 33, 5466–5474 (2013).

19. Favila, S. E., Chanales, A. J. H. & Kuhl, B. A. Experience-dependent hippocampal pattern differentiation prevents interference during subsequent learning. Nat. Commun. 7, 11066 (2016).

20. Chanales, A. J. H., Oza, A., Favila, S. E. & Kuhl, B. A. Overlap among Spatial Memories Triggers Repulsion of Hippocampal Representations. Curr. Biol. CB 27, 2307-2317.e5 (2017).

21. Schapiro, A. C., Kustner, L. V. & Turk-Browne, N. B. Shaping of Object Representations in the Human Medial Temporal Lobe Based on Temporal Regularities. Curr. Biol. 22, 1622– 1627 (2012).

22. Kim, G., Norman, K. A. & Turk-Browne, N. B. Neural differentiation of incorrectly predicted memories. J. Neurosci. (2017) doi:10.1523/JNEUROSCI.3272-16.2017.

23. Jiang, J., Wang, S.-F., Guo, W., Fernandez, C. & Wagner, A. D. Prefrontal reinstatement of contextual task demand is predicted by separable hippocampal patterns. Nat. Commun. 11, 2053 (2020).

24. Hulbert, J. C. & Norman, K. A. Neural Differentiation Tracks Improved Recall of Competing Memories Following Interleaved Study and Retrieval Practice. Cereb. Cortex N. Y. N 1991 25, 3994–4008 (2015).

25. Tompary, A. & Davachi, L. Consolidation Promotes the Emergence of Representational Overlap in the Hippocampus and Medial Prefrontal Cortex. Neuron 96, 228-241.e5 (2017).

26. Schapiro, A. C., Rogers, T. T., Cordova, N. I., Turk-Browne, N. B. & Botvinick, M. M. Neural representations of events arise from temporal community structure. Nat. Neurosci. 16, 486–492 (2013).

27. Schapiro, A. C., Turk-Browne, N. B., Norman, K. A. & Botvinick, M. M. Statistical learning of temporal community structure in the hippocampus. Hippocampus 26, 3–8 (2016).

28. Schapiro, A. C., Gregory, E., Landau, B., McCloskey, M. & Turk-Browne, N.B. The Necessity of the Medial Temporal Lobe for Statistical Learning. J. Cogn. Neurosci. 26, 1736– 1747 (2014).

29. Zeithamova, D., Dominick, A. L. & Preston, A. R. Hippocampal and Ventral Medial Prefrontal Activation during Retrieval-Mediated Learning Supports Novel Inference. Neuron 75, 168–179 (2012).

30. Vaidya, A. R., Jones, H. M., Castillo, J. & Badre, D. Neural representation of abstract task structure during generalization. eLife 10, e63226 (2021).

31. Kumaran, D. & McClelland, J. L. Generalization through the recurrent interaction of episodic memories: a model of the hippocampal system. Psychol. Rev. 119, 573–616 (2012).

32. Kumaran, D., Hassabis, D. & McClelland, J. L. What Learning Systems do Intelligent Agents Need? Complementary Learning Systems Theory Updated. Trends Cogn. Sci. 20, 512–534 (2016).

33. Schapiro, A. C., Turk-Browne, N. B., Botvinick, M. M. & Norman, K. A. Complementary learning systems within the hippocampus: a neural network modelling approach to reconciling episodic memory with statistical learning. Philos. Trans. R. Soc. B Biol. Sci. 372, 20160049 (2017).

34. Gupta, A. S., van der Meer, M. A. A., Touretzky, D. S. & Redish, A. D. Hippocampal replay is not a simple function of experience. Neuron 65, 695–705 (2010).

35. Wu, X. & Foster, D. J. Hippocampal Replay Captures the Unique Topological Structure of a Novel Environment. J. Neurosci. 34, 6459–6469 (2014).

36. Shohamy, D. & Wagner, A. D. Integrating Memories in the Human Brain: Hippocampal-Midbrain Encoding of Overlapping Events. Neuron 60, 378–389 (2008).

37. Ólafsdóttir, H. F., Bush, D. & Barry, C. The Role of Hippocampal Replay in Memory and Planning. Curr. Biol. 28, R37–R50 (2018).

38. Yu, J. Y., Liu, D. F., Loback, A., Grossrubatscher, I. & Frank, L. M. Specific hippocampal representations are linked to generalized cortical representations in memory. Nat. Commun. 9, 2209 (2018).

39. Jadhav, S. P., Rothschild, G., Roumis, D. K. & Frank, L. M. Coordinated Excitation and Inhibition of Prefrontal Ensembles during Awake Hippocampal Sharp-Wave Ripple Events. Neuron 90, 113–127 (2016).

40. Lewis, P. A. & Durrant, S. J. Overlapping memory replay during sleep builds cognitive schemata. Trends Cogn. Sci. 15, 343–351 (2011).

41. Kuhl, B. A., Bainbridge, W. A. & Chun, M. M. Neural Reactivation Reveals Mechanisms for Updating Memory. J. Neurosci. 32, 3453–3461 (2012).

42. Tse, D. et al. Schemas and memory consolidation. Science 316, 76–82 (2007).

43. Tse, D. et al. Schema-Dependent Gene Activation and Memory Encoding in Neocortex. Science 333, 891–895 (2011).

44. Ranganath, C., Heller, A., Cohen, M. X., Brozinsky, C. J. & Rissman, J. Functional connectivity with the hippocampus during successful memory formation. Hippocampus 15, 997–1005 (2005).

45. Siapas, A. G., Lubenov, E. V. & Wilson, M. A. Prefrontal phase locking to hippocampal theta oscillations. Neuron 46, 141–151 (2005).

46. Preston, A. R. & Eichenbaum, H. Interplay of hippocampus and prefrontal cortex in memory. Curr. Biol. CB 23, R764–773 (2013).

47. Kumaran, D., Summerfield, J. J., Hassabis, D. & Maguire, E. A. Tracking the Emergence of Conceptual Knowledge during Human Decision Making. Neuron 63, 889–901 (2009).

48. Zeithamova, D. & Preston, A. R. Flexible memories: Differential roles for medial temporal lobe and prefrontal cortex in cross-episode binding. J. Neurosci. 30, 14676–14684 (2010).

49. Frankland, P. W. & Bontempi, B. The organization of recent and remote memories. Nat. Rev. Neurosci. 6, 119–130 (2005).

50. Takashima, A. et al. Declarative memory consolidation in humans: a prospective functional magnetic resonance imaging study. Proc. Natl. Acad. Sci. U. S. A. 103, 756–761 (2006).

51. Takehara-Nishiuchi, K. & McNaughton, B. L. Spontaneous changes of neocortical code for associative memory during consolidation. Science 322, 960–963 (2008).

52. Zheng, L., Gao, Z., McAvan, A. S., Isham, E. A. & Ekstrom, A. D. Partially overlapping spatial environments trigger reinstatement in hippocampus and schema representations in prefrontal cortex. Nat. Commun. 12, 6231 (2021).

53. Chrastil, E. R. Neural evidence supports a novel framework for spatial navigation. Psychon. Bull. Rev. 20, 208–227 (2013).

54. O’Keefe, J. & Dostrovsky, J. The hippocampus as a spatial map. Preliminary evidence from unit activity in the freely-moving rat. Brain Res. 34, 171–175 (1971).

55. McNaughton, B. L., Battaglia, F. P., Jensen, O., Moser, E. I. & Moser, M.-B. Path integration and the neural basis of the ‘cognitive map’. Nat. Rev. Neurosci. 7, 663–678 (2006).

56. Hafting, T., Fyhn, M., Molden, S., Moser, M.-B. & Moser, E. I. Microstructure of a spatial map in the entorhinal cortex. Nature 436, 801–806 (2005).

57. Kropff, E., Carmichael, J. E., Moser, M.-B. & Moser, E. I. Speed cells in the medial entorhinal cortex. Nature 523, 419–424 (2015).

58. Deuker, L., Bellmund, J. L., Navarro Schröder, T. & Doeller, C. F. An event map of memory space in the hippocampus. eLife 5, e16534 (2016).

59. Peer, M., Brunec, I. K., Newcombe, N. S. & Epstein, R. A. Structuring Knowledge with Cognitive Maps and Cognitive Graphs. Trends Cogn. Sci. 25, 37–54 (2021).

60. Nyberg, N., Duvelle, É., Barry, C. & Spiers, H. J. Spatial goal coding in the hippocampal formation. Neuron 110, 394–422 (2022).

61. Howard, L. R. et al. The hippocampus and entorhinal cortex encode the path and Euclidean distances to goals during navigation. Curr. Biol. CB 24, 1331–1340 (2014).

62. Patai, E. Z. et al. Hippocampal and Retrosplenial Goal Distance Coding After Long-term Consolidation of a Real-World Environment. Cereb. Cortex 29, 2748–2758 (2019).

63. Weisberg, S. M., Schinazi, V. R., Newcombe, N. S., Shipley, T. F. & Epstein, R. A. Variations in cognitive maps: understanding individual differences in navigation. J. Exp. Psychol. Learn. Mem. Cogn. 40, 669–682 (2014).

64. Weisberg, S. M. & Newcombe, N. S. How do (some) people make a cognitive map? Routes, places, and working memory. J. Exp. Psychol. Learn. Mem. Cogn. 42, 768–785 (2016).

65. Weisberg, S. M. & Newcombe, N. S. Cognitive Maps: Some People Make Them, Some People Struggle. Curr. Dir. Psychol. Sci. 27, 220–226 (2018).

66. Brown, T. I., Ross, R. S., Keller, J. B., Hasselmo, M. E. & Stern, C. E. Which way was I going? Contextual retrieval supports the disambiguation of well learned overlapping navigational routes. J. Neurosci. Off. J. Soc. Neurosci. 30, 7414–7422 (2010).

67. McKenzie, S. et al. Hippocampal Representation of Related and Opposing Memories Develop within Distinct, Hierarchically Organized Neural Schemas. Neuron 83, 202–215 (2014).

68. Milivojevic, B. & Doeller, C. F. Mnemonic networks in the hippocampal formation: from spatial maps to temporal and conceptual codes. J. Exp. Psychol. Gen. 142, 1231–1241 (2013).

69. Chrastil, E. R. & Warren, W. H. From Cognitive Maps to Cognitive Graphs. PLOS ONE 9, e112544 (2014).

70. Wolbers, T. & Hegarty, M. What determines our navigational abilities? Trends Cogn. Sci. 14, 138–146 (2010).

71. Solway, A., Miller, J. F. & Kahana, M. J. PandaEPL: A library for programming spatial navigation experiments. Behav. Res. Methods 45, (2013).

72. Clarke, A., Pell, P. J., Ranganath, C. & Tyler, L. K. Learning Warps Object Representations in the Ventral Temporal Cortex. J. Cogn. Neurosci. 28, 1010–1023 (2016).

73. Olsen, R. K. et al. Performance-Related Sustained and Anticipatory Activity in Human Medial Temporal Lobe during Delayed Match-to-Sample. J. Neurosci. 29, 11880–11890 (2009).

74. Chang, L. J. et al. Endogenous variation in ventromedial prefrontal cortex state dynamics during naturalistic viewing reflects affective experience. Sci. Adv. 7, eabf7129.

75. Walther, A. et al. Reliability of dissimilarity measures for multi-voxel pattern analysis. NeuroImage 137, 188–200 (2016).

76. Bates, D., Mächler, M., Bolker, B. & Walker, S. Fitting Linear Mixed-Effects Models Using lme4. J. Stat. Softw. 67, 1–48 (2015).

77. Kuznetsova, A., Brockhoff, P. B. & Christensen, R. H. B. lmerTest Package: Tests in Linear Mixed Effects Models. J. Stat. Softw. 82, 1–26 (2017).

78. Ritchey, M., Wing, E. A., LaBar, K. S. & Cabeza, R. Neural similarity between encoding and retrieval is related to memory via hippocampal interactions. Cereb. Cortex N. Y. N 1991 23, 2818–2828 (2013).

