## Supplementary Materials for "Representational integration and differentiation in the human hippocampus following goal-directed navigation"

### Methods

*Stimuli.* Nineteen building facades were selected from https://www.lughertexture.com and used to create landmarks in the virtual environment. Facades were selected to have a variety of features and be of different architectural styles and colors. After resizing, removing signs and reflections, and adding roofs in Adobe Photoshop (Adobe, Inc.), buildings were rendered onto 3D models of equal shape and size. Twelve buildings were randomly selected to be incorporated into each participant's unique environment. Fractal images marked goal locations within the virtual environment. Twenty-three fractals were drawn from the Dryad Digital Repository[1,2] and from the stimulus set used in Brown et al., 2016[3]. Images were resized and rendered onto 3D models of equal shape and size. Fifteen fractals were randomly selected to be incorporated into each participant's unique environment. Each participant's unique set of building and fractal stimuli were used in both behavioral navigation tasks and all fMRI sessions. Buildings and fractals were matched for low-level visual features by using the SHINE toolbox in Matlab (version R2015b) to equate their luminance histograms[4].

All 3D models for the virtual environment were created in Maya (Autodesk, Inc.), a 3D graphics program. Textures were acquired via a Google search and https://www.lughertexture.com. While the specific stimuli used for landmarks varied between participants, the positions of the landmarks were fixed in the environment. However, goal locations were selected to fall within a 3 arbitrary unit (a.u.) radius of each landmark (1 a.u. = ~3 m/10 ft); thus, goal locations differed slightly across participants.

*Anatomical data preprocessing.* fMRI data preprocessing was performed with fMRIPrep 1.5.3rc2 (RRID:SCR_016216)[5,6], which is based on *Nipype* 1.3.1 (RRID:SCR_002502) [7,8]. The T1-weighted (T1w) structural images were corrected for intensity non-uniformity (INU) with N4BiasFieldCorrection[9], distributed with ANTs 2.2.0 (RRID:SCR_004757)[10], and used as the T1w-reference throughout the workflow. The T1w-reference was then skull-stripped with a *Nipype* implementation of the antsBrainExtraction.sh workflow (from ANTs), using OASIS30ANTs as the target template. Brain tissue segmentation of cerebrospinal fluid (CSF), white-matter (WM) and gray-matter (GM) was performed on the brain-extracted T1w-reference using FAST (FSL 5.0.9, RRID:SCR_002823)[11]. Brain surfaces were reconstructed using recon-all (FreeSurfer 6.0.1, RRID:SCR_001847)[12], and the brain mask estimated previously was refined with a custom variation of the method to reconcile ANTs-derived and FreeSurfer-derived segmentations of the cortical gray-matter of Mindboggle (RRID:SCR_002438)[13]. Volume-based spatial normalization to standardized space (MNI152NLin2009cAsym) was performed through nonlinear registration with antsRegistration (ANTs 2.2.0), using brain-extracted versions of both the T1w-reference and T1w-template. The ICBM 152 Nonlinear Asymmetrical template version 2009 [RRID:SCR_008796; TemplateFlow ID:MNI152NLin2009cAsym][14] was selected for spatial normalization.

*Functional data preprocessing.* For each of the 12 functional runs per participant (4 per scan session), the following preprocessing was performed. First, a reference volume and its skull-stripped version were generated using a custom methodology of fMRIPrep. A B0-nonuniformity map (or fieldmap) was estimated based on the two EPI references with opposing phase-encoding directions, with 3dQwarp (AFNI)[15]. Based on the estimated susceptibility distortion, a corrected EPI reference was calculated for a more accurate co-registration with the anatomical reference. The BOLD reference was then co-registered to the T1w-reference using bbregister (FreeSurfer) which implements boundary-based registration[16]. Co-registration was configured with six degrees of freedom. Head-motion parameters with respect to the BOLD reference (transformation matrices and six corresponding rotation and translation parameters) were estimated before any spatiotemporal filtering using MCFLIRT (FSL 5.0.9)[17]. The BOLD time-series were resampled onto their original, native space by applying a single, composite transform to correct for head-motion and susceptibility distortions. These resampled BOLD time-series will be referred to as preprocessed BOLD in original space, or just preprocessed BOLD. The BOLD time-series were resampled into standard space, generating a preprocessed BOLD run in MNI space. Several confounding time-series were calculated based on the preprocessed BOLD: framewise displacement (FD), DVARS and a set of low-frequency regressors for temporal high-pass filtering. FD and DVARS were calculated for each functional run, both using their implementations in

Nipype (following the definitions by Power et al., 2014[18]). The head-motion estimates calculated in the correction step were also placed within the corresponding confounds file. All resamplings were performed with a single interpolation step by composing all the pertinent transformations (i.e. head-motion transform matrices, susceptibility distortion correction when available, and co-registrations to anatomical and output spaces). Gridded (volumetric) resamplings were performed using antsApplyTransforms (ANTs), configured with Lanczos interpolation to minimize the smoothing effects of other kernels[19]. The preprocessed fMRI data were smoothed by a 2-mm full-width-half-maximum Gaussian kernel.

*Linear Mixed-Effects Models*. To control for perceptual similarity between task stimuli, we input the stimuli into vNet, a deep neural network model. vNet was trained on ecoset, a large-scale image set containing images from 565 basic level categories[20]. We extracted hidden layer representations putatively corresponding to V1 and IT cortex (layers 1 and 10, respectively), computed similarity matrices, and included these estimates as regressors in linear mixed effects models. Open-source code is available at https://codeocean.com/capsule/9570390/tree/v1).

For all mixed-effects models, the standard set of random effects was chosen by taking a maximal model with all random effects indicated by the experiment setup, then incrementally removing effects and testing the nested model fits using the likelihood ratio test. This procedure was performed for all ROIs. The final standard set of random effects was composed of the minimum necessary effects to achieve the best fit in all regions, and included a random intercept modeling the mean subject-specific outcome value, as well as random slope terms modeling subject-specific effects of independent variables of interest (i.e., scan session and average univariate activation).

In all mixed-effects models, scan session was dummy-coded with Pre-Learning (Day 1) as the baseline. Stimulus type and context were sum-coded with the contrasts of landmark > fractal and same track > different tracks, respectively. Behavioral performance group was sum-coded with the contrast ME > LE.

Models were estimated using a restricted maximum likelihood (REML) approach. Model convergence issues were resolved by changing from the default optimizer to "bobyqa" and not having the model calculate derivatives.

### Results

**Local and global distance in entorhinal cortex.** As with the hippocampus, we tested whether pattern similarity for landmark buildings in EC scaled with local (within-track) and global (across-track) distance. An initial model of local distance that included hemisphere (left and right EC) and link distance (1 and 2) revealed interactions between scan session and hemisphere (Day 2 > Day 1 X hemisphere, $\beta = 0.050 \pm 0.023$, $t = 2.093$, $p = 0.036$; $d = 0.47$; Day 3 > Day 1 X hemisphere, $\beta = 0.069 \pm 0.024$, $t = 2.835$, $p = 0.005$, survived FDR correction; $d = 0.67$; Supplementary Table 14). However, in contrast to hippocampus, we found no interactions between distance and scan session (Supplementary Table 15-16) when we fit models to data from each hemisphere individually. An initial model of global distance that included hemisphere (left and right EC) and link distance (2, 3, and 4) revealed no main effect of hemisphere or interactions with hemisphere (Supplementary Table 17). Unlike the hippocampus, we found no interactions between distance and scan session in the global model (Supplementary Table 18). Our findings in EC were not surprising, as the extent to which EC itself can support structured representations is unclear. Spatial properties of EC neurons are known to be important for path integration and the building of structured knowledge in the hippocampus and neocortex[21,22]. Yet, hippocampal conjunctive coding and interactions between the hippocampus, EC, and neocortex are necessary for cross event-generalization and retrieval-mediated learning[23–27].

**Control analyses.** All linear mixed-effects models were fit to data from a visual control region defined as a conjunction of FreeSurfer's pericalcarine and calcarine sulcus regions in both hemispheres.

We first tested whether context effects differed between the visual control region, the hippocampus, and EC by running a complete model predicting neural pattern similarity, with scan session (Pre-Learning/Day 1, Post Local Navigation/Day 2, and

Post Global Navigation/Day 3), context (same path and different paths), and region (calcarine, hippocampus, and EC) as predictors. Region was dummy-coded with the visual control region (calcarine) serving as the baseline. Here we found main effects (hippocampus: $\beta$ = -0.44 ± 0.003, t = -148.995, p < 2e$^{-16}$, survived FDR correction, d = -32.51; EC: $\beta$ = -0.458 ± 0.003, t = -156.298, p < 2e$^{-16}$, survived FDR correction, d = -34.10) and region X scan session interactions for all regions (hippocampus X Day 3 > Day 1, $\beta$ = 0.008 ± 0.004, t = 2.202, p = 0.027, d = 0.48; EC X Day 2 > Day 1, $\beta$ = -0.008 ± 0.004, t = -2.019, p = 0.043, d = -0.42; EC X Day 3 > Day 1, $\beta$ = 0.017 ± 0.004, t = 4.354, p < 1.34e$^{-5}$, d = 0.95; Supplementary Table 30).

We then fit a linear mixed-effects model predicting neural pattern similarity to data from the visual control region. Scan session (Pre-Learning/Day 1, Post Local Navigation/Day 2, and Post Global Navigation/Day 3), stimulus type (landmarks and fractals), and context (same track and different tracks) were included as predictors. When the context model was fit to data from the control region, we found no interactions between context and scan session (Day 2 > Day 1 X context, $\beta$ = -0.003 ± 0.005; t = -0.717; p = 0.474; Day 3 > Day 1 X context, $\beta$ = -0.002 ± 0.005; t = -0.416; p = 0.678) or context, stimulus type, and scan session (Day 2 > Day 1 X stimulus type X context, $\beta$ = -0.004 ± 0.009; t = -0.377; p = 0.706; Day 3 > Day 1 X stimulus type X context, $\beta$ = 0.014 ± 0.01; t = 1.509; p = 0.131; Supplementary Fig. 4; Supplementary Table 31).

We next tested whether distance effects differed between the visual control region, the hippocampus, and vmPFC by running a complete model predicting neural pattern similarity for landmarks, with scan session (Pre-Learning/Day 1, Post Local Navigation/Day 2, and Post Global Navigation/Day 3), link distance, and region (calcarine, hippocampus, and vmPFC) as predictors. Region was dummy-coded with the visual control region (calcarine) serving as the baseline. Here we found main effects (hippocampus: $\beta$ = -0.506 ± 0.011, t = -46.889, p < 2e$^{-16}$, survived FDR correction, d = -9.78; vmPFC: $\beta$ = -0.483 ± 0.013, t = -38.294, p < 2e$^{-16}$, survived FDR correction, d = -7.98) and region X scan session interactions for both regions (hippocampus X Day 2 > Day 1, $\beta$ = 0.077 ± 0.015, t = 5.102, p < 3.38e$^{-7}$, d = 1.06, hippocampus X Day 3 > Day 1, $\beta$ = 0.047 ± 0.015, t = 3.079, p = 0.002, d = 0.67; vmPFC X Day 2 > Day 1, $\beta$ = 0.089 ± 0.017, t = 5.129, p < 2.92e$^{-7}$, d = 1.07; Supplementary Table 26).

We fit a linear mixed-effects model predicting neural pattern similarity between landmarks on the same track to data from the visual control region. Scan session (Pre-Learning/Day 1, Post Local Navigation/Day 2, and Post Global Navigation/Day 3) and link distance (1 and 2) were included as predictors. When the local distance model was fit to data from the control region, we found no interactions between link distance and scan session (Day 2 > Day 1 X distance, $\beta = 0.007 \pm 0.022$; $t = 0.304$; $p = 0.761$; Day 3 > Day 1 X distance, $\beta = -0.035 \pm 0.022$; $t = -1.57$; $p = 0.117$, Supplementary Fig. 5a; Supplementary Table 32). Finally, we fit a linear mixed-effects model predicting neural pattern similarity between landmarks on different tracks to data from the visual control region. Scan session (Pre-Learning/Day 1, Post Local Navigation/Day 2, and Post Global Navigation/Day 3) and link distance (2, 3, and 4) were included as predictors. When this distance model was fit to data from the control region, we found no interactions between link distance and scan session (Day 2 > Day 1 X distance, $\beta = -0.003 \pm 0.01$; $t = -0.261$; $p > 0.794$; Day 3 > Day 1 X distance, $\beta = 0.007 \pm 0.01$; $t = 0.727$; $p > 0.468$; Supplementary Fig. 5b; Supplementary Table 33). These control analyses indicate that the effects observed in the hippocampus were not observed throughout the brain.

### References


1. Wilming, N. *et al.* An extensive dataset of eye movements during viewing of complex images. *Sci. Data* **4**, 160126 (2017).
2. Wilming, N. *et al.* Data from: An extensive dataset of eye movements during viewing of complex images. 7495233822 bytes (2017) doi:10.5061/DRYAD.9PF75.
3. Brown, T. I. *et al.* Prospective representation of navigational goals in the human hippocampus. *Science* **352**, 1323–1326 (2016).
4. Willenbockel, V. *et al.* Controlling low-level image properties: The SHINE toolbox. *Behav. Res. Methods* **42**, 671–684 (2010).
5. Esteban, O. *et al.* fMRIPrep: a robust preprocessing pipeline for functional MRI. *Nat. Methods* **16**, 111–116 (2019).

6. Esteban, O. *et al. fMRIPrep: a robust preprocessing pipeline for functional MRI*. (Zenodo, 2022). doi:10.5281/zenodo.5898602.
7. Gorgolewski, K. *et al.* Nipype: A Flexible, Lightweight and Extensible Neuroimaging Data Processing Framework in Python. *Front. Neuroinformatics* **5**, (2011).
8. Esteban, O. *et al. nipy/nipype: 1.7.1*. (Zenodo, 2022). doi:10.5281/zenodo.6415183.
9. Tustison, N. J. *et al.* N4ITK: improved N3 bias correction. *IEEE Trans. Med. Imaging* **29**, 1310–1320 (2010).
10. Avants, B. B., Epstein, C. L., Grossman, M. & Gee, J. C. Symmetric diffeomorphic image registration with cross-correlation: evaluating automated labeling of elderly and neurodegenerative brain. *Med. Image Anal.* **12**, 26–41 (2008).
11. Zhang, Y., Brady, M. & Smith, S. Segmentation of brain MR images through a hidden Markov random field model and the expectation-maximization algorithm. *IEEE Trans. Med. Imaging* **20**, 45–57 (2001).
12. Dale, A. M., Fischl, B. & Sereno, M. I. Cortical surface-based analysis. I. Segmentation and surface reconstruction. *NeuroImage* **9**, 179–194 (1999).
13. Klein, A. *et al.* Mindboggling morphometry of human brains. *PLOS Comput. Biol.* **13**, e1005350 (2017).
14. Fonov, V., Evans, A., McKinstry, R., Almli, C. & Collins, D. Unbiased nonlinear average age-appropriate brain templates from birth to adulthood. *NeuroImage* **47**, S102 (2009).
15. Cox, R. W. & Hyde, J. S. Software tools for analysis and visualization of fMRI data. *NMR Biomed.* **10**, 171–178 (1997).
16. Greve, D. N. & Fischl, B. Accurate and robust brain image alignment using boundary-based registration. *NeuroImage* **48**, 63–72 (2009).
17. Jenkinson, M., Bannister, P., Brady, M. & Smith, S. Improved Optimization for the Robust and Accurate Linear Registration and Motion Correction of Brain Images. *NeuroImage* **17**, 825–841 (2002).

18. Power, J. D. *et al.* Methods to detect, characterize, and remove motion artifact in resting state fMRI. *NeuroImage* **84**, 320–341 (2014).
19. Lanczos, C. Evaluation of Noisy Data. *J. Soc. Ind. Appl. Math. Ser. B Numer. Anal.* **1**, 76–85 (1964).
20. Mehrer, J., Spoerer, C. J., Jones, E. C., Kriegeskorte, N. & Kietzmann, T. C. An ecologically motivated image dataset for deep learning yields better models of human vision. *Proc. Natl. Acad. Sci. U. S. A.* **118**, e2011417118 (2021).
21. McNaughton, B. L., Battaglia, F. P., Jensen, O., Moser, E. I. & Moser, M.-B. Path integration and the neural basis of the 'cognitive map'. *Nat. Rev. Neurosci.* **7**, 663–678 (2006).
22. Hafting, T., Fyhn, M., Molden, S., Moser, M.-B. & Moser, E. I. Microstructure of a spatial map in the entorhinal cortex. *Nature* **436**, 801–806 (2005).
23. Kumaran, D. & McClelland, J. L. Generalization through the recurrent interaction of episodic memories: a model of the hippocampal system. *Psychol. Rev.* **119**, 573–616 (2012).
24. Kumaran, D., Hassabis, D. & McClelland, J. L. What Learning Systems do Intelligent Agents Need? Complementary Learning Systems Theory Updated. *Trends Cogn. Sci.* **20**, 512–534 (2016).
25. Zeithamova, D., Dominick, A. L. & Preston, A. R. Hippocampal and Ventral Medial Prefrontal Activation during Retrieval-Mediated Learning Supports Novel Inference. *Neuron* **75**, 168–179 (2012).
26. Preston, A. R. & Eichenbaum, H. Interplay of hippocampus and prefrontal cortex in memory. *Curr. Biol. CB* **23**, R764-773 (2013).
27. Schapiro, A. C., Turk-Browne, N. B., Botvinick, M. M. & Norman, K. A. Complementary learning systems within the hippocampus: a neural network modelling approach to reconciling episodic memory with statistical learning. *Philos. Trans. R. Soc. B Biol. Sci.* **372**, 20160049 (2017).

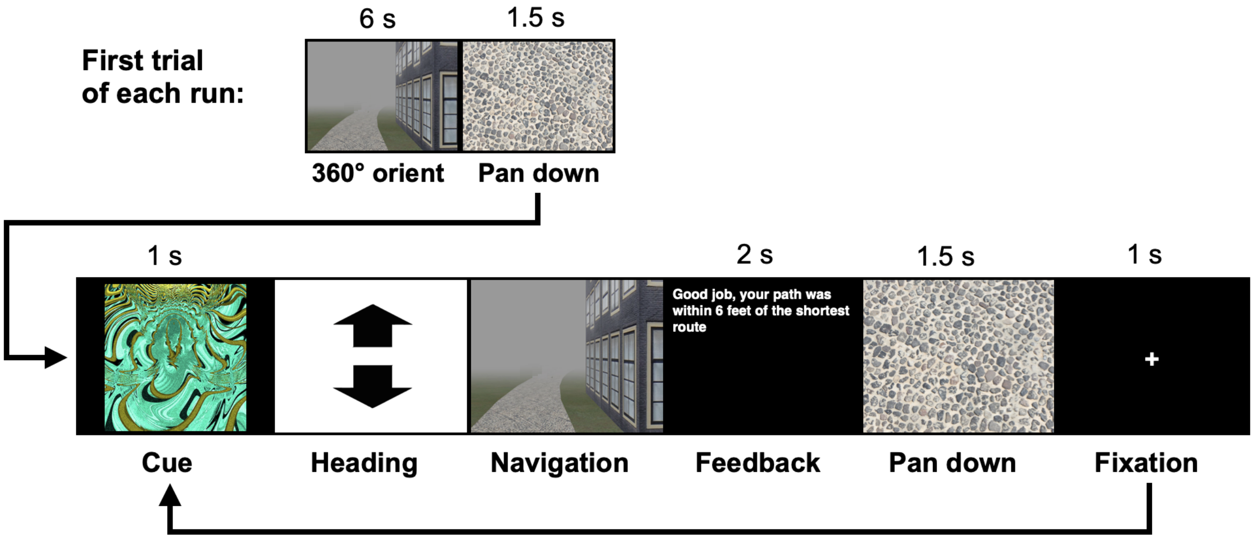


**Supplementary Figure 1**. Trial structure for all navigation tasks. Each behavioral run contained 10 navigation trials. At the start of each run, participants were placed at a location on the track and rotated 360 degrees (6 s). Trials then proceeded as follows: 1) a fractal cue appeared onscreen (1 s) indicating the goal to which the participant should navigate; 2) participants chose their heading direction and 3) navigated to the cued goal location, pressing the spacebar when they arrived; 4) feedback appeared onscreen revealing whether the participant was at the correct location and whether they had navigated via the shortest path (2 s); and 5) the camera panned down and a fixation cross appeared (1 s) before the next trial began. On learning trials, goal locations were marked by fractal images appearing on the track. On test trials, fractals were not visible on the track and participants had to rely on memory to navigate.

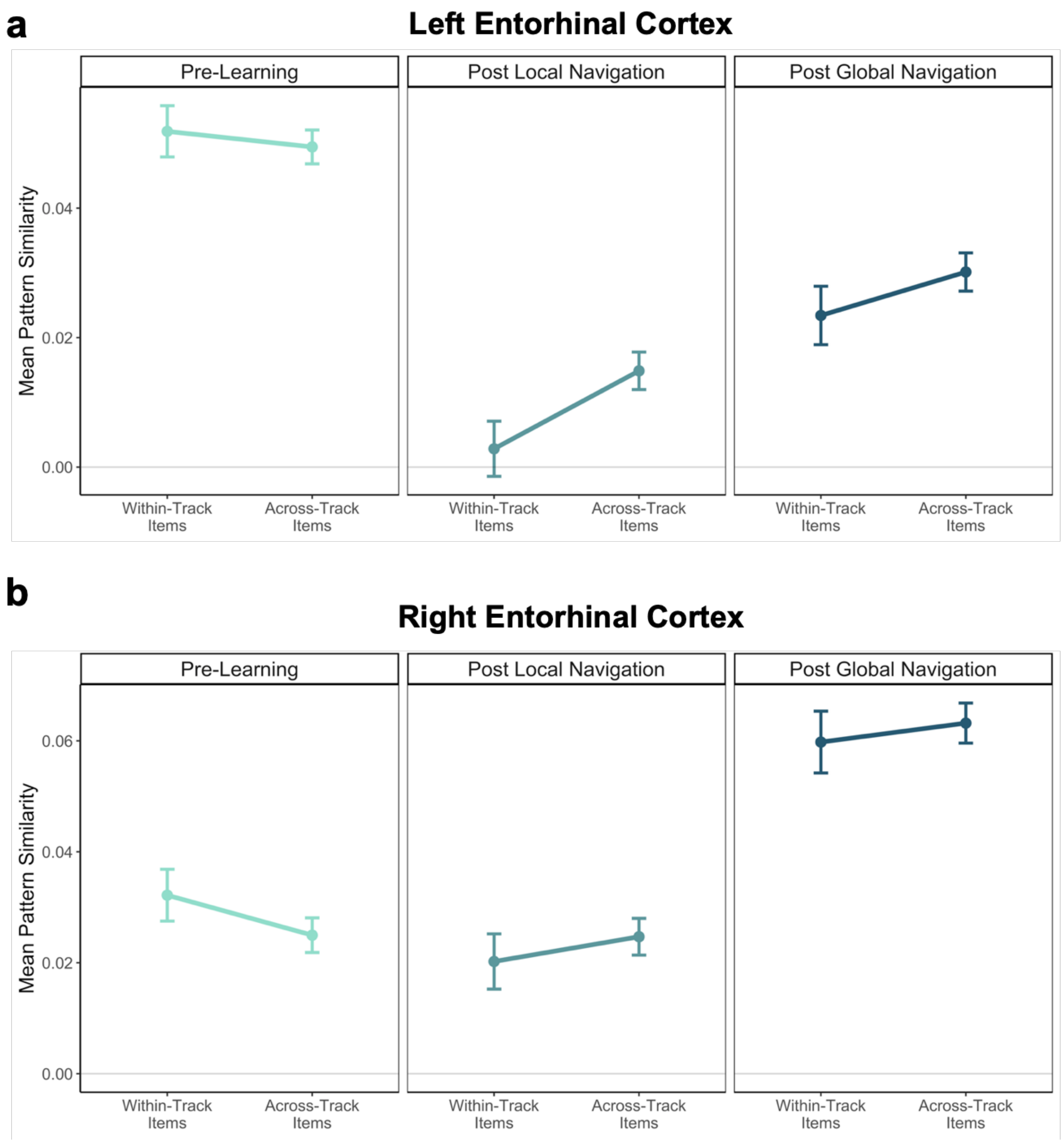


**Supplementary Figure 2**. Pattern similarity for items located on the same vs. different tracks in EC. (A) Mean pattern similarity in Left EC. (B) Mean pattern similarity in Right EC. The experiment was conducted once (Error bars denote SE of the estimates. Left EC, n = 20 biologically independent samples; Right EC, n = 18 biologically independent samples). Source data are provided as a source data file.

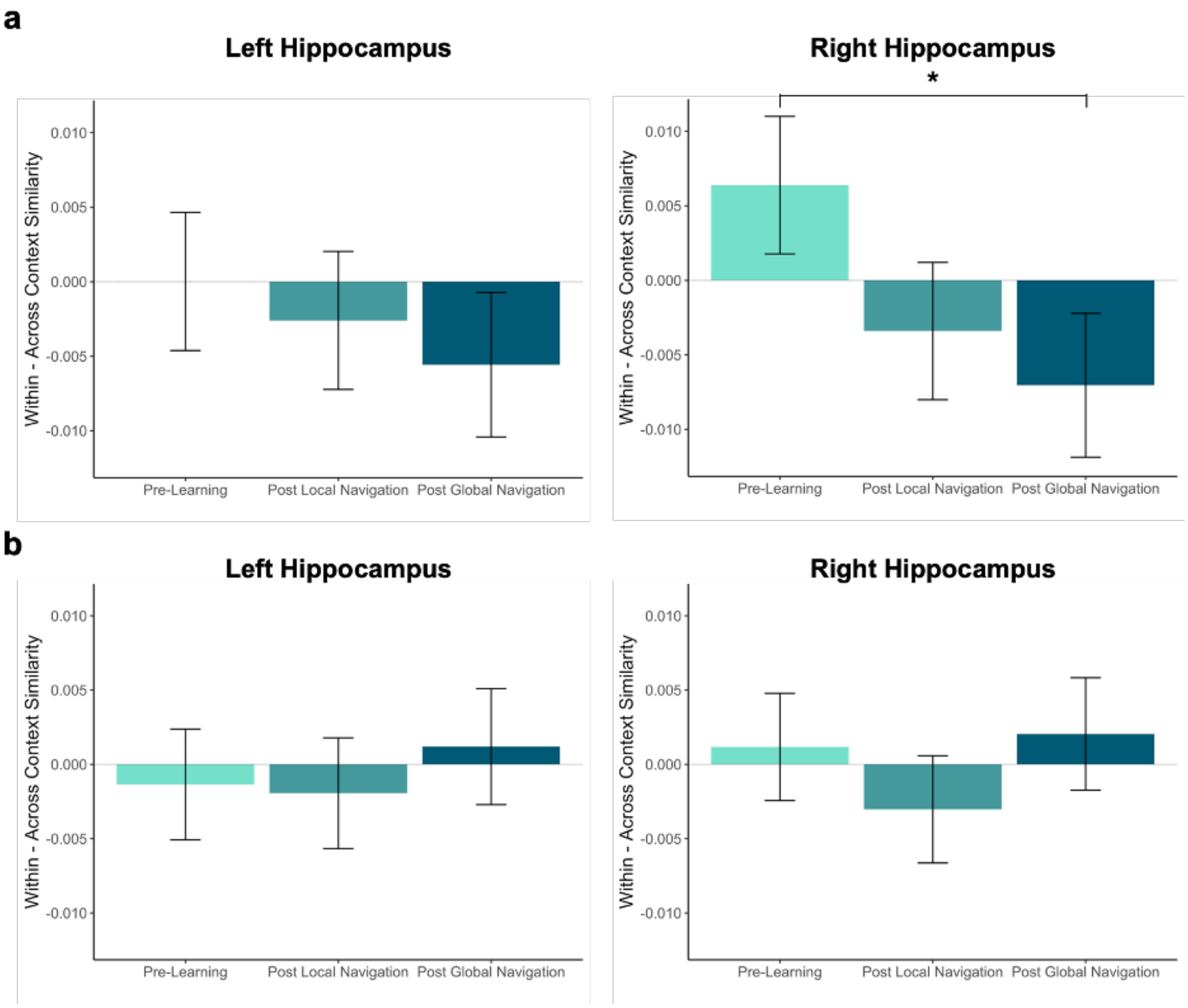


**Supplementary Figure 3**. Contrast estimates for models fit to data in the left and right hippocampus. (A) Within - across context similarity for landmark buildings. Scan X context interaction for D3>D1 ($\beta$ = -0.01; t = -2.014; $p < 0.045$; d = -0.44) (B) Within - across context similarity for fractals. (* $p < 0.05$. Error bars denote SE of the estimates. Day 2 > Day 1, n = 23; Day 3 > Day 1, n = 21; examined within a single experiment. Source data are provided as a source data file).

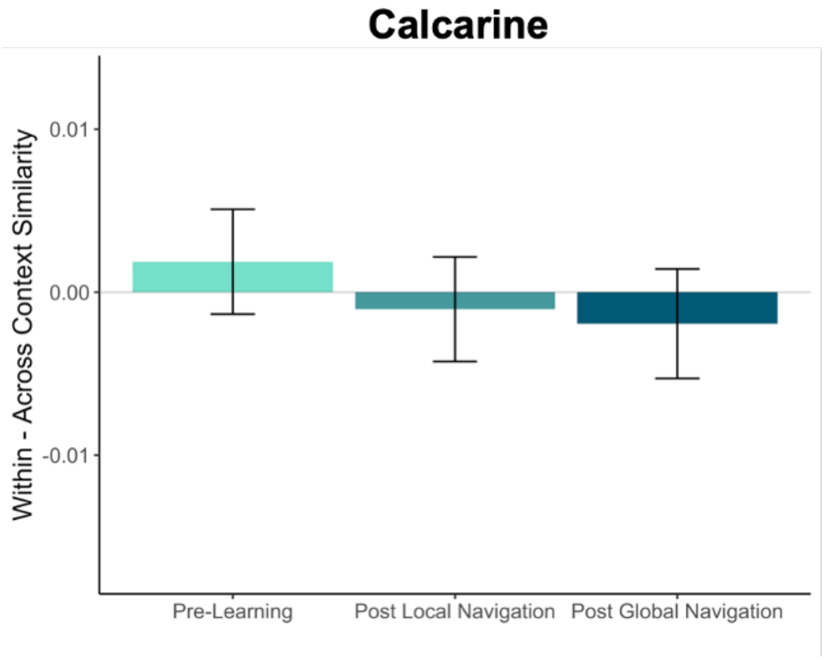


**Supplementary Figure 4**. Contrast estimates for context models fit to data in a visual region serving as a control (calcarine). Within - across context similarity for landmark buildings. Interactions between scan session and context were not significant. (Error bars denote SE of the estimates. Day 2 > Day 1, n = 23; Day 3 > Day 1, n = 21; examined within a single experiment. Source data are provided as a source data file).

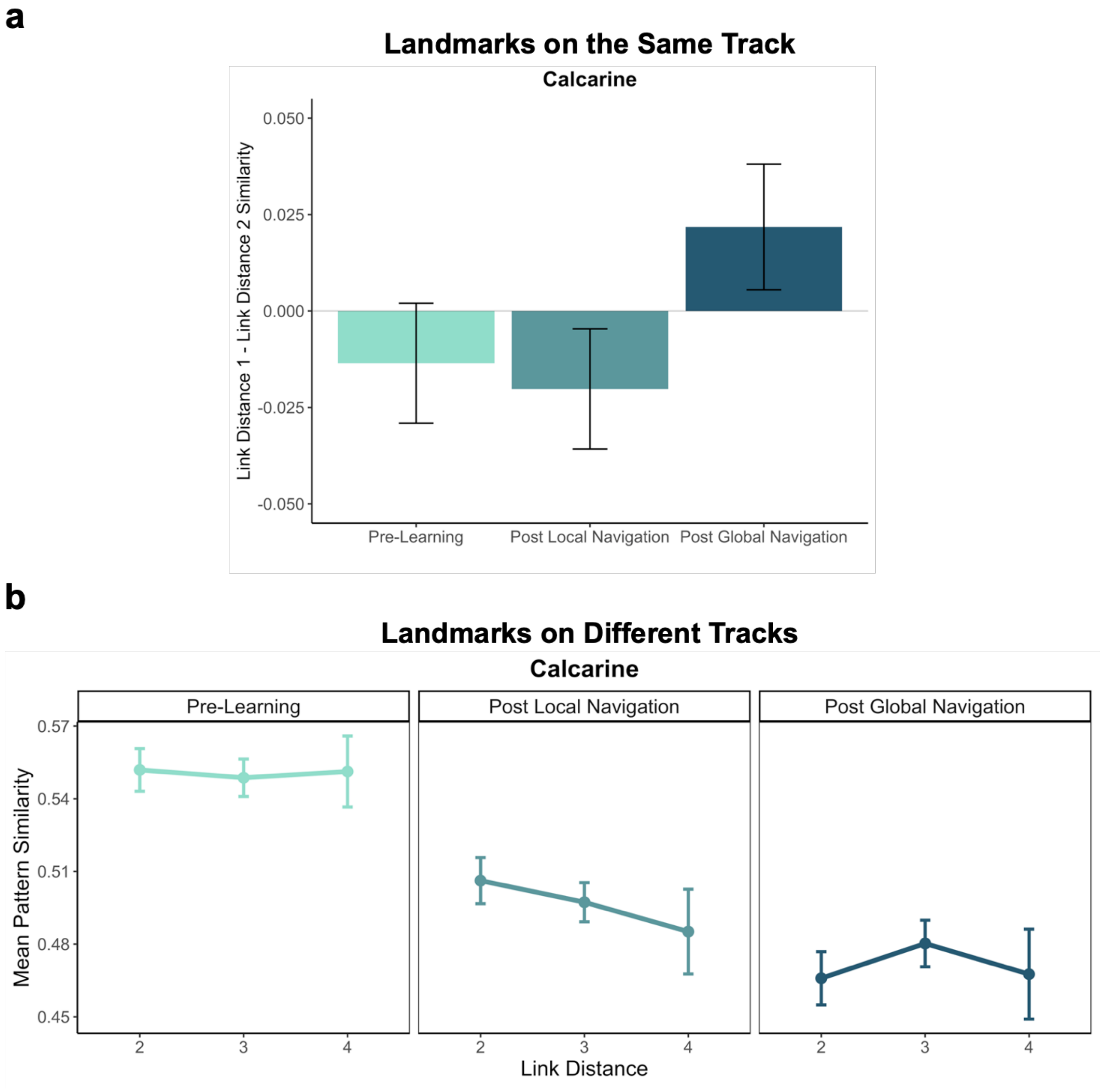


**Supplementary Figure 5**. Pattern similarity for landmark buildings at different link distances in a visual region serving as a control (calcarine). (A) Model contrast estimates for landmarks on the same track following the Local and Global Navigation Tasks. (B) Pattern similarity for landmarks on different tracks Pre-Learning (left), after the Local Navigation Task (center), and after the Global Navigation Task (right). Interactions between link distance and scan session were not significant. (Error bars denote SE of the estimates. Day 2 > Day 1, n = 23; Day 3 > Day 1, n = 21; examined within a single experiment. Source data are provided as a source data file).

**a**

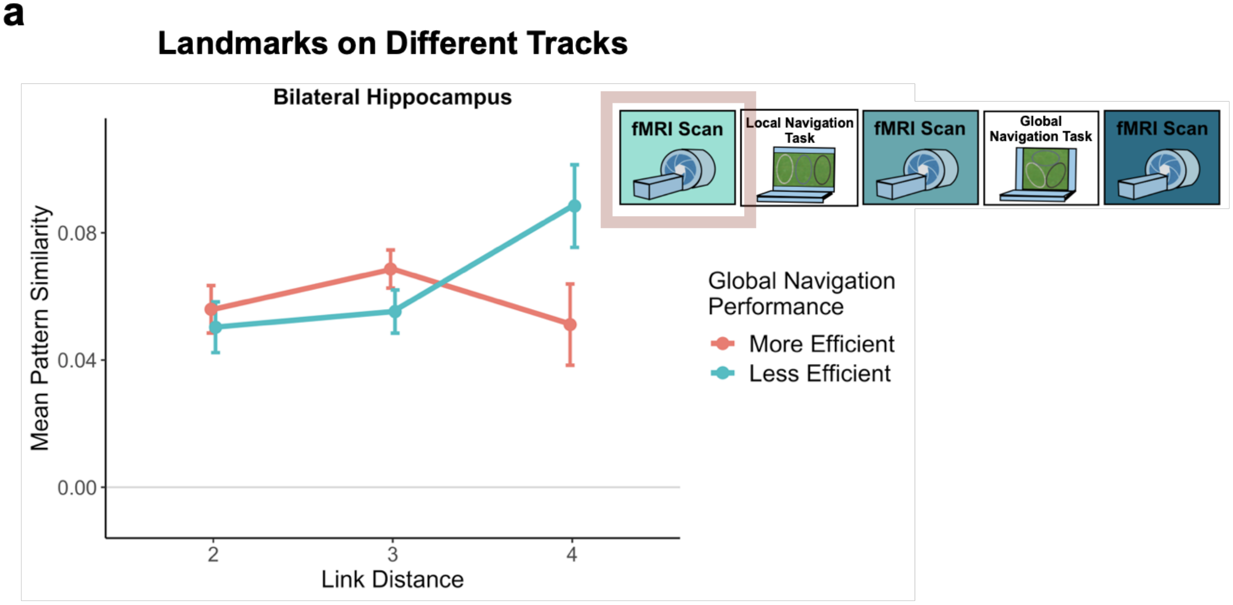


**b**

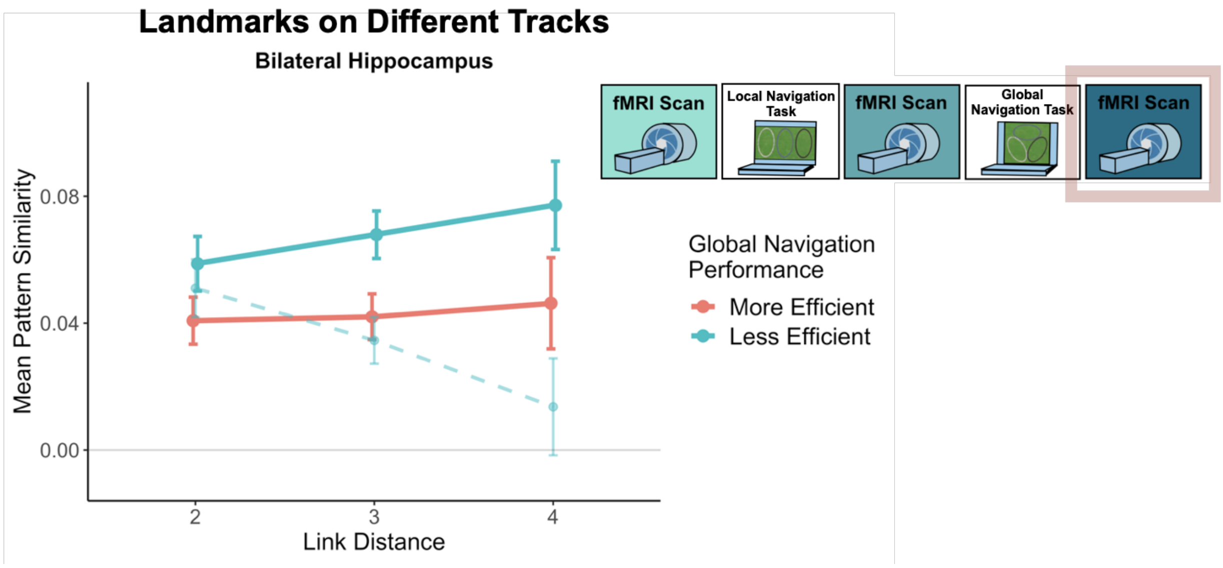


**Supplementary Figure 6**. Hippocampal pattern similarity for landmark buildings on different tracks. Pattern similarity relationships did not differ between participants who are more or less efficient on the Global Navigation Task, (A) Pre-Learning (Day 1) and (B) Post Global Navigation (Day 3). Dashed line represents pattern similarity for Less Efficient participants Post Local Navigation. Data are split by performance as shown in Fig. 5a. (Error bars denote SE of the estimates. More Efficient, n = 11; Less Efficient, n = 10; examined within a single experiment. Source data are provided as a source data file).

a

b

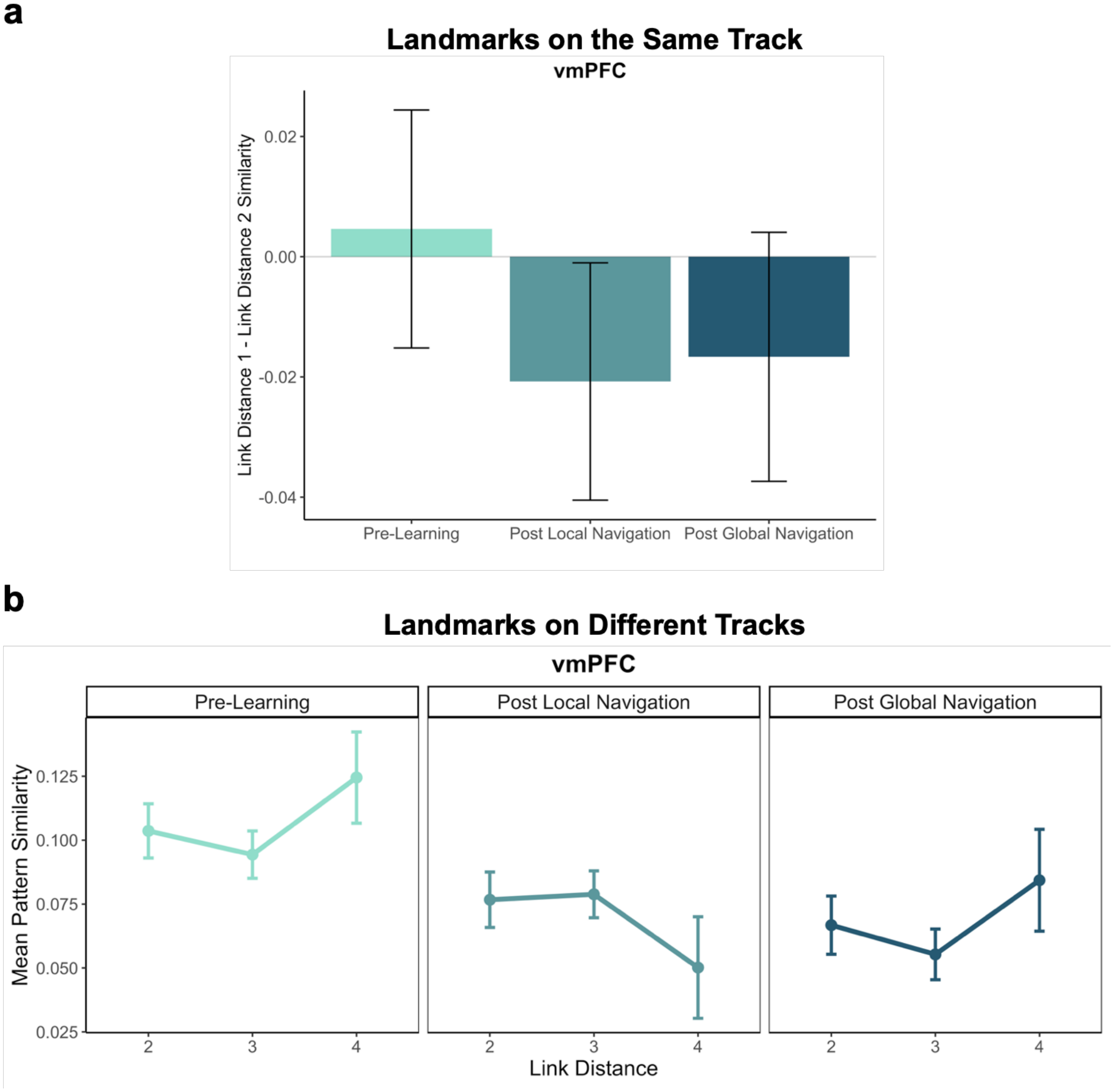


**Supplementary Figure 7**. Pattern similarity for landmark buildings at different link distances in vmPFC. (A) Model contrast estimates for landmarks on the same track following the Local and Global Navigation Tasks. (B) vmPFC pattern similarity for landmarks on different tracks Pre-Learning (left), after the Local Navigation Task (center), and after the Global Navigation Task (right). Interactions between link distance and scan session were not significant. (Error bars denote SE of the estimates. Day 2 > Day 1, n = 23; Day 3 > Day 1, n = 21; examined within a single experiment. Source data are provided as a source data file).

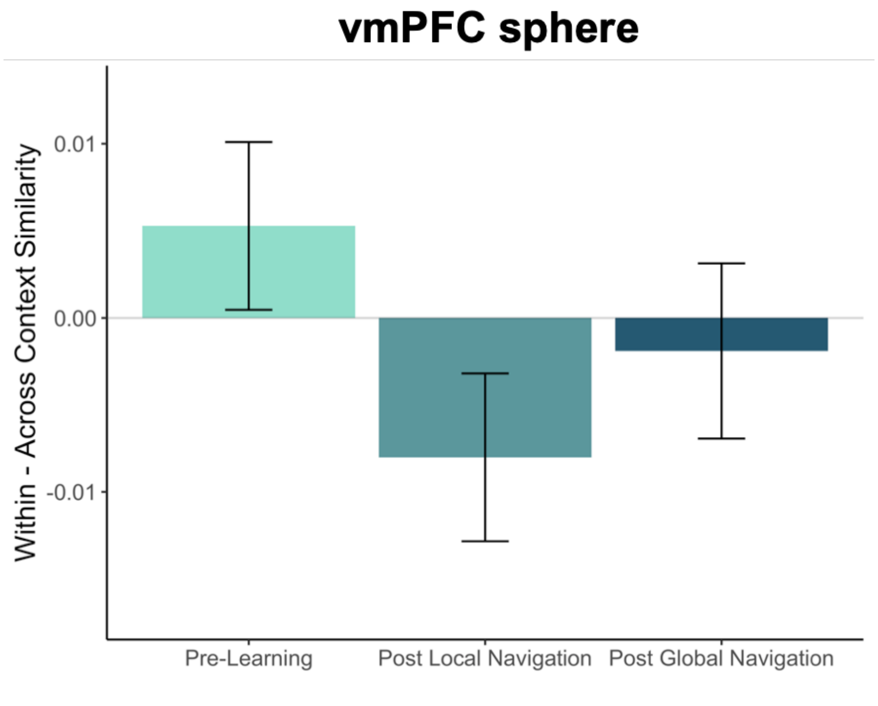


**Supplementary Figure 8**. Contrast estimates for context models fit to data from an 8mm-radius spherical ROI in vmPFC. Within - across context similarity for landmark buildings. Interactions between scan session and context were not significant. (Error bars denote SE of the estimates. Day 2 > Day 1, n = 23; Day 3 > Day 1, n = 21; examined within a single experiment. Source data are provided as a source data file).

**Supplementary Table 1.** Linear mixed-effects model results for an omnibus context model predicting neural pattern similarity, fit to data in EC. Source data are provided as a Source Data file. SE = standard error. No correction for multiple comparisons was applied.

| **Variable** | ***β*** | **SE** | ***t*** | ***p*** |
|---|---|---|---|---|
| (Intercept) | 0.049 | 0.017 | 2.912 | **0.004** |
| Day 2 | -0.037 | 0.020 | -1.874 | 0.061 |
| Day 3 | -0.023 | 0.020 | -1.178 | 0.239 |
| stimulus type (landmark > fractal) | 0.008 | 0.005 | 1.607 | 0.108 |
| context (same track > different tracks) | 0.002 | 0.005 | 0.461 | 0.645 |
| hemisphere | -0.022 | 0.004 | -6.060 | **$1.37e^{-9}$** |
| Day 2 X stimulus type | 0.010 | 0.007 | 1.457 | 0.145 |
| Day 3 X stimulus type | -0.003 | 0.007 | -0.358 | 0.720 |
| Day 2 X context | -0.014 | 0.007 | -2.046 | **0.041** |
| Day 3 X context | -0.010 | 0.007 | -1.394 | 0.163 |
| stimulus type X context | -0.001 | 0.010 | -0.139 | 0.889 |
| Day 2 X hemisphere | 0.022 | 0.005 | 4.192 | **$2.77e^{-5}$** |
| Day 3 X hemisphere | 0.056 | 0.005 | 10.552 | **$<2e^{-16}$** |
| stimulus type X hemisphere | 0.000 | 0.007 | 0.036 | 0.971 |

| | | | | |
|---|---|---|---|---|
| context X hemisphere | 0.004 | 0.007 | 0.524 | 0.600 |
| Day 2 X stimulus type X context | 0.002 | 0.014 | 0.132 | 0.895 |
| Day 3 X stimulus type X context | -0.005 | 0.014 | -0.387 | 0.699 |
| Day 2 X stimulus type X hemisphere | -0.011 | 0.010 | -1.132 | 0.258 |
| Day 3 X stimulus type X hemisphere | -0.003 | 0.010 | -0.319 | 0.749 |
| Day 2 X context X hemisphere | 0.003 | 0.010 | 0.263 | 0.793 |
| Day 3 X context X hemisphere | 0.001 | 0.010 | 0.117 | 0.907 |
| stimulus type X context X hemisphere | -0.009 | 0.014 | -0.633 | 0.527 |
| Day 2 X stimulus type X context X hemisphere | -0.001 | 0.020 | -0.050 | 0.960 |
| Day 3 X stimulus type X context X hemisphere | 0.022 | 0.020 | 1.057 | 0.291 |

**Supplementary Table 2.** Linear mixed-effects model results for a context model predicting neural pattern similarity, fit to data in left EC. Source data are provided as a Source Data file. SE = standard error. No correction for multiple comparisons was applied.

| **Variable** | ***β*** | **SE** | ***t*** | ***p*** |
|---|---|---|---|---|
| (Intercept) | 0.082 | 0.040 | 2.025 | **0.043** |
| Day 2 | -0.036 | 0.024 | -1.490 | 0.136 |
| Day 3 | -0.026 | 0.021 | -1.223 | 0.221 |

| | | | | |
|---|---|---|---|---|
| context (same track > different tracks) | 0.003 | 0.004 | 0.656 | 0.512 |
| Day 2 X context | -0.014 | 0.006 | -2.339 | **0.019** |
| Day 3 X context | -0.009 | 0.006 | -1.457 | 0.145 |

**Supplementary Table 3.** Linear mixed-effects model results for a context model predicting neural pattern similarity, fit to data in right EC. Source data are provided as a Source Data file. SE = standard error. No correction for multiple comparisons was applied.

| **Variable** | ***β*** | **SE** | ***t*** | ***p*** |
|---|---|---|---|---|
| (Intercept) | 0.067 | 0.049 | 1.362 | 0.173 |
| Day 2 | 0.002 | 0.022 | 0.071 | 0.943 |
| Day 3 | 0.044 | 0.024 | 1.836 | 0.066 |
| context (same track > different tracks) | 0.008 | 0.005 | 1.455 | 0.146 |
| Day 2 X context | -0.012 | 0.007 | -1.568 | 0.117 |
| Day 3 X context | -0.011 | 0.008 | -1.404 | 0.160 |

**Supplementary Table 4.** Linear mixed-effects model results for an omnibus context model predicting neural pattern similarity, fit to data in the hippocampus. Source data are provided as a Source Data file. SE = standard error. No correction for multiple comparisons was applied.

| **Variable** | ***β*** | **SE** | ***t*** | ***p*** |
|---|---|---|---|---|
| (Intercept) | 0.035 | 0.009 | 3.858 | **$8.11e^{-4}$** |
| Day 2 | 0.002 | 0.011 | 0.136 | 0.892 |

| | | | | |
|---|---|---|---|---|
| Day 3 | -0.006 | 0.012 | -0.542 | 0.588 |
| stimulus type (landmark > fractal) | 0.007 | 0.003 | 2.422 | **0.015** |
| context (same track > different tracks) | -0.001 | 0.003 | -0.254 | 0.800 |
| hemisphere | 0.006 | 0.002 | 2.756 | **0.006** |
| Day 2 X stimulus type | -0.003 | 0.004 | -0.715 | 0.475 |
| Day 3 X stimulus type | -0.011 | 0.004 | -2.604 | **0.009** |
| Day 2 X context | -0.002 | 0.004 | -0.365 | 0.715 |
| Day 3 X context | -0.001 | 0.004 | -0.334 | 0.739 |
| stimulus type X context | 0.001 | 0.006 | 0.246 | 0.806 |
| Day 2 X hemisphere | -0.011 | 0.003 | -3.477 | **$5.08e^{-4}$** |
| Day 3 X hemisphere | 0.012 | 0.003 | 3.795 | **$1.48e^{-4}$** |
| stimulus type X hemisphere | 0.004 | 0.004 | 0.866 | 0.387 |
| context X hemisphere | 0.005 | 0.004 | 1.066 | 0.287 |
| Day 2 X stimulus type X context | -0.002 | 0.009 | -0.216 | 0.829 |
| Day 3 X stimulus type X context | -0.008 | 0.009 | -0.920 | 0.357 |
| Day 2 X stimulus type X hemisphere | -0.002 | 0.006 | -0.320 | 0.749 |
| Day 3 X stimulus type X hemisphere | -0.001 | 0.006 | -0.192 | 0.848 |

| | | | | |
|---|---|---|---|---|
| Day 2 X context X hemisphere | -0.005 | 0.006 | -0.899 | 0.369 |
| Day 3 X context X hemisphere | -0.005 | 0.006 | -0.767 | 0.443 |
| stimulus type X context X hemisphere | 0.004 | 0.009 | 0.497 | 0.619 |
| Day 2 X stimulus type X context X hemisphere | -0.004 | 0.012 | -0.322 | 0.747 |
| Day 3 X stimulus type X context X hemisphere | -0.006 | 0.012 | -0.511 | 0.610 |

**Supplementary Table 5.** Linear mixed-effects model results for a context model predicting neural pattern similarity for landmark buildings, fit to data in left hippocampus. Source data are provided as a Source Data file. SE = standard error. No correction for multiple comparisons was applied.

| **Variable** | ***β*** | **SE** | ***t*** | ***p*** |
|---|---|---|---|---|
| (Intercept) | 0.051 | 0.049 | 1.043 | 0.297 |
| Day 2 | 0.001 | 0.014 | 0.058 | 0.954 |
| Day 3 | -0.011 | 0.016 | -0.686 | 0.493 |
| context (same track > different tracks) | 0.000 | 0.005 | 0.003 | 0.998 |
| Day 2 X context | -0.003 | 0.007 | -0.400 | 0.689 |
| Day 3 X context | -0.006 | 0.007 | -0.834 | 0.404 |

**Supplementary Table 6.** Linear mixed-effects model results for a context model predicting neural pattern similarity for landmark buildings, fit to data in right hippocampus. Source data are provided as a Source Data file. SE = standard error. No correction for multiple comparisons was applied.

| Variable | *β* | SE | *t* | *p* |
|---|---|---|---|---|
| (Intercept) | 0.062 | 0.049 | 1.260 | 0.208 |
| Day 2 | -0.011 | 0.015 | -0.747 | 0.455 |
| Day 3 | -0.002 | 0.017 | -0.099 | 0.921 |
| context (same track > different tracks) | 0.006 | 0.005 | 1.385 | 0.166 |
| Day 2 X context | -0.010 | 0.007 | -1.502 | 0.133 |
| Day 3 X context | -0.013 | 0.007 | -2.014 | **0.044** |

**Supplementary Table 7.** Linear mixed-effects model results for a context model predicting neural pattern similarity for fractals, fit to data in left hippocampus. Source data are provided as a Source Data file. SE = standard error. No correction for multiple comparisons was applied.

| Variable | *β* | SE | *t* | *p* |
|---|---|---|---|---|
| (Intercept) | 0.029 | 0.030 | 0.945 | 0.345 |
| Day 2 | 0.003 | 0.012 | 0.262 | 0.794 |
| Day 3 | -0.002 | 0.011 | -0.150 | 0.881 |
| context (same track > different tracks) | -0.001 | 0.004 | -0.363 | 0.717 |
| Day 2 X context | -0.001 | 0.005 | -0.111 | 0.911 |
| Day 3 X context | 0.003 | 0.005 | 0.472 | 0.637 |

**Supplementary Table 8.** Linear mixed-effects model results for a context model predicting neural pattern similarity for fractals, fit to data in right hippocampus. Source data are provided as a Source Data file. SE = standard error. No correction for multiple comparisons was applied.

| **Variable** | ***β*** | **SE** | ***t*** | ***p*** |
|---|---|---|---|---|
| (Intercept) | 0.100 | 0.029 | 3.398 | **6.89e$^{-4}$** |
| Day 2 | -0.003 | 0.012 | -0.268 | 0.789 |
| Day 3 | 0.011 | 0.014 | 0.817 | 0.414 |
| context (same track > different tracks) | 0.001 | 0.004 | 0.326 | 0.745 |
| Day 2 X context | -0.004 | 0.005 | -0.823 | 0.411 |
| Day 3 X context | 0.001 | 0.005 | 0.166 | 0.868 |

**Supplementary Table 9.** Linear mixed-effects model results for a context model predicting neural pattern similarity that included EC and hippocampus. Source data are provided as a Source Data file. SE = standard error. No correction for multiple comparisons was applied.

| **Variable** | ***β*** | **SE** | ***t*** | ***p*** |
|---|---|---|---|---|
| (Intercept) | 0.074 | 0.021 | 3.623 | **3.66e$^{-4}$** |
| Day 2 | -0.012 | 0.013 | -0.883 | 0.377 |
| Day 3 | -0.010 | 0.013 | -0.780 | 0.436 |
| context (same track > different tracks) | 0.001 | 0.003 | 0.473 | 0.636 |
| region | -0.015 | 0.002 | -7.360 | **1.85e$^{-13}$** |

| | | | | |
|---|---|---|---|---|
| Day 2 X context | -0.004 | 0.004 | -0.956 | 0.339 |
| Day 3 X context | -0.002 | 0.004 | -0.594 | 0.553 |
| Day 2 X region | -0.017 | 0.003 | -5.757 | **8.58e$^{-9}$** |
| Day 3 X region | 0.014 | 0.003 | 4.592 | **4.40e$^{-6}$** |
| context X region | 0.004 | 0.004 | 0.907 | 0.364 |
| Day 2 X context X region | -0.009 | 0.006 | -1.635 | 0.102 |
| Day 3 X context X region | -0.007 | 0.006 | -1.278 | 0.201 |

**Supplementary Table 10.** Linear mixed-effects model results for an omnibus model of local (within-track) distance, predicting neural pattern similarity in the hippocampus. Source data are provided as a Source Data file. SE = standard error. No correction for multiple comparisons was applied.

| **Variable** | ***β*** | **SE** | ***t*** | ***p*** |
|---|---|---|---|---|
| (Intercept) | 0.044 | 0.012 | 3.621 | **9.12e$^{-4}$** |
| Day 2 | -0.016 | 0.015 | -1.041 | 0.298 |
| Day 3 | -0.020 | 0.016 | -1.236 | 0.217 |
| distance (link distance 2 > link distance 1) | -0.014 | 0.015 | -0.974 | 0.330 |
| hemisphere | 0.006 | 0.011 | 0.528 | 0.598 |
| Day 2 X distance | 0.036 | 0.021 | 1.721 | 0.085 |

| | | | | |
|---|---|---|---|---|
| Day 3 X distance | 0.033 | 0.021 | 1.520 | 0.129 |
| Day 2 X hemisphere | -0.011 | 0.015 | -0.723 | 0.470 |
| Day 3 X hemisphere | 0.004 | 0.015 | 0.288 | 0.774 |
| distance X hemisphere | -0.003 | 0.021 | -0.124 | 0.901 |
| Day 2 X distance X hemisphere | 0.008 | 0.030 | 0.280 | 0.780 |
| Day 3 X distance X hemisphere | 0.004 | 0.030 | 0.121 | 0.904 |

**Supplementary Table 11.** Linear mixed-effects model results in a model of local (within-track) distance, predicting neural pattern similarity in the hippocampus. Source data are provided as a Source Data file. SE = standard error. No correction for multiple comparisons was applied.

| **Variable** | ***β*** | **SE** | ***t*** | ***p*** |
|---|---|---|---|---|
| (Intercept) | -0.037 | 0.088 | -0.419 | 0.675 |
| Day 2 | -0.021 | 0.013 | -1.597 | 0.110 |
| Day 3 | -0.018 | 0.014 | -1.247 | 0.213 |
| distance (link distance 2 > link distance 1) | -0.016 | 0.010 | -1.510 | 0.131 |
| Day 2 X distance | 0.040 | 0.015 | 2.700 | **0.007** |
| Day 3 X distance | 0.034 | 0.015 | 2.259 | **0.024** |

**Supplementary Table 12.** Linear mixed-effects model results in an omnibus model of global (across-track) distance, predicting neural pattern similarity in the hippocampus. Source data are provided as a Source Data file. SE = standard error. No correction for multiple comparisons was applied.

| Variable | *β* | SE | *t* | *p* |
|---|---|---|---|---|
| (Intercept) | 0.033 | 0.016 | 2.023 | **0.043** |
| Day 2 | 0.018 | 0.023 | 0.779 | 0.436 |
| Day 3 | -0.010 | 0.023 | -0.445 | 0.657 |
| distance | 0.002 | 0.007 | 0.369 | 0.712 |
| hemisphere | -0.012 | 0.018 | -0.659 | 0.510 |
| Day 2 X distance | -0.012 | 0.010 | -1.297 | 0.195 |
| Day 3 X distance | 0.000 | 0.010 | 0.044 | 0.965 |
| Day 2 X hemisphere | 0.019 | 0.025 | 0.763 | 0.445 |
| Day 3 X hemisphere | 0.016 | 0.026 | 0.632 | 0.527 |
| distance X hemisphere | 0.011 | 0.010 | 1.159 | 0.246 |
| Day 2 X distance X hemisphere | -0.015 | 0.013 | -1.082 | 0.279 |
| Day 3 X distance X hemisphere | -0.007 | 0.014 | -0.485 | 0.628 |

**Supplementary Table 13.** Linear mixed-effects model results in a model of global (across-track) distance, predicting neural pattern similarity in the hippocampus. Source data are provided as a Source Data file. SE = standard error. No correction for multiple comparisons was applied.

| Variable | *β* | SE | *t* | *p* |
|---|---|---|---|---|

| | | | | |
|---|---|---|---|---|
| (Intercept) | 0.076 | 0.064 | 1.181 | 0.238 |
| Day 2 | 0.057 | 0.027 | 2.126 | **0.034** |
| Day 3 | 0.002 | 0.027 | 0.077 | 0.939 |
| distance | 0.007 | 0.005 | 1.567 | 0.117 |
| Day 2 X distance | -0.020 | 0.007 | -2.891 | **0.004** |
| Day 3 X distance | -0.003 | 0.007 | -0.413 | 0.679 |

**Supplementary Table 14.** Linear mixed-effects model results in an omnibus model of local (within-track) distance, predicting neural pattern similarity in EC. Source data are provided as a Source Data file. SE = standard error. No correction for multiple comparisons was applied.

| **Variable** | ***β*** | **SE** | ***t*** | ***p*** |
|---|---|---|---|---|
| (Intercept) | 0.053 | 0.021 | 2.521 | **0.012** |
| Day 2 | -0.057 | 0.023 | -2.503 | **0.012** |
| Day 3 | -0.040 | 0.024 | -1.662 | 0.097 |
| distance (link distance 2 > link distance 1) | 0.011 | 0.024 | 0.475 | 0.635 |
| hemisphere | -0.030 | 0.017 | -1.775 | 0.076 |
| Day 2 X distance | 0.006 | 0.033 | 0.191 | 0.849 |
| Day 3 X distance | 0.008 | 0.034 | 0.226 | 0.821 |

| | | | | |
|---|---|---|---|---|
| Day 2 X hemisphere | 0.050 | 0.024 | 2.093 | **0.036** |
| Day 3 X hemisphere | 0.069 | 0.024 | 2.835 | **0.005** |
| distance X hemisphere | -0.034 | 0.033 | -1.031 | 0.303 |
| Day 2 X distance X hemisphere | 0.029 | 0.047 | 0.620 | 0.535 |
| Day 3 X distance X hemisphere | 0.026 | 0.048 | 0.545 | 0.586 |

**Supplementary Table 15.** Linear mixed-effects model results in a model of local (within-track) distance, predicting neural pattern similarity in left EC. Source data are provided as a Source Data file. SE = standard error. No correction for multiple comparisons was applied.

| **Variable** | ***β*** | **SE** | ***t*** | ***p*** |
|---|---|---|---|---|
| (Intercept) | 0.109 | 0.180 | 0.604 | 0.546 |
| Day 2 | -0.057 | 0.022 | -2.529 | **0.012** |
| Day 3 | -0.043 | 0.023 | -1.920 | 0.055 |
| distance (link distance 2 > link distance 1) | 0.011 | 0.021 | 0.508 | 0.611 |
| Day 2 X distance | 0.007 | 0.030 | 0.238 | 0.812 |
| Day 3 X distance | 0.007 | 0.031 | 0.232 | 0.816 |

**Supplementary Table 16.** Linear mixed-effects model results in a model of local (within-track) distance, predicting neural pattern similarity in right EC. Source data are provided as a Source Data file. SE = standard error. No correction for multiple comparisons was applied.

| Variable | *β* | SE | *t* | *p* |
|---|---|---|---|---|
| (Intercept) | -0.410 | 0.213 | -1.921 | 0.055 |
| Day 2 | -0.001 | 0.024 | -0.060 | 0.952 |
| Day 3 | 0.029 | 0.028 | 1.056 | 0.291 |
| distance (link distance 2 > link distance 1) | -0.025 | 0.025 | -0.977 | 0.329 |
| Day 2 X distance | 0.034 | 0.036 | 0.937 | 0.349 |
| Day 3 X distance | 0.038 | 0.037 | 1.043 | 0.297 |

**Supplementary Table 17.** Linear mixed-effects model results in an omnibus model of global (across-track) distance, predicting neural pattern similarity in EC. Source data are provided as a Source Data file. SE = standard error. No correction for multiple comparisons was applied.

| Variable | *β* | SE | *t* | *p* |
|---|---|---|---|---|
| (Intercept) | 0.062 | 0.026 | 2.393 | **0.017** |
| Day 2 | 0.003 | 0.036 | 0.091 | 0.928 |
| Day 3 | -0.036 | 0.035 | -1.042 | 0.297 |
| distance | 0.002 | 0.011 | 0.194 | 0.846 |
| hemisphere | -0.031 | 0.028 | -1.096 | 0.273 |
| Day 2 X distance | -0.024 | 0.015 | -1.555 | 0.120 |

| | | | | |
|---|---|---|---|---|
| Day 3 X distance | 0.001 | 0.016 | 0.084 | 0.933 |
| Day 2 X hemisphere | 0.008 | 0.040 | 0.208 | 0.835 |
| Day 3 X hemisphere | 0.055 | 0.041 | 1.336 | 0.182 |
| distance X hemisphere | 0.001 | 0.015 | 0.069 | 0.945 |
| Day 2 X distance X hemisphere | 0.013 | 0.021 | 0.614 | 0.540 |
| Day 3 X distance X hemisphere | 0.003 | 0.022 | 0.139 | 0.889 |

**Supplementary Table 18.** Linear mixed-effects model results in a model of global (across-track) distance, predicting neural pattern similarity in EC. Source data are provided as a Source Data file. SE = standard error. No correction for multiple comparisons was applied.

| **Variable** | ***β*** | **SE** | ***t*** | ***p*** |
|---|---|---|---|---|
| (Intercept) | 0.139 | 0.102 | 1.361 | 0.174 |
| Day 2 | 0.034 | 0.042 | 0.812 | 0.417 |
| Day 3 | -0.014 | 0.041 | -0.328 | 0.743 |
| distance | 0.002 | 0.008 | 0.201 | 0.841 |
| Day 2 X distance | -0.017 | 0.011 | -1.607 | 0.108 |
| Day 3 X distance | 0.003 | 0.011 | 0.260 | 0.795 |

**Supplementary Table 19.** Linear mixed-effects model results from a model predicting hippocampal pattern similarity Post Local Navigation (Day 2) for the two performance groups. Source data are provided as a Source Data file. SE = standard error. No correction for multiple comparisons was applied.

| Variable | *β* | SE | *t* | *p* |
|---|---|---|---|---|
| (Intercept) | 0.072 | 0.117 | 0.617 | 0.537 |
| distance | -0.004 | 0.005 | -0.687 | 0.492 |
| performance group (more efficient > less efficient) | -0.056 | 0.042 | -1.342 | 0.180 |
| distance X performance group | 0.025 | 0.011 | 2.386 | **0.017** |

**Supplementary Table 20.** Linear mixed-effects model results from a model predicting hippocampal pattern similarity Pre-Learning (Day 1) for the two performance groups. Source data are provided as a Source Data file. SE = standard error. No correction for multiple comparisons was applied.

| Variable | *β* | SE | *t* | *p* |
|---|---|---|---|---|
| (Intercept) | 0.037 | 0.101 | 0.366 | 0.714 |
| distance | 0.008 | 0.005 | 1.820 | 0.069 |
| performance group (more efficient > less efficient) | 0.036 | 0.037 | 0.978 | 0.328 |
| distance X performance group | -0.004 | 0.009 | -0.460 | 0.645 |

**Supplementary Table 21.** Linear mixed-effects model results from a model predicting hippocampal pattern similarity Post Global Navigation (Day 3) for the two performance groups. Source data are provided as a Source Data file. SE = standard error. No correction for multiple comparisons was applied.

| Variable | *β* | SE | *t* | *p* |
|---|---|---|---|---|

| | | | | |
|---|---|---|---|---|
| (Intercept) | 0.116 | 0.112 | 1.035 | 0.301 |
| distance | 0.005 | 0.005 | 0.941 | 0.347 |
| performance group (more efficient > less efficient) | 0.010 | 0.042 | 0.248 | 0.804 |
| distance X performance group | -0.005 | 0.010 | -0.472 | 0.637 |

**Supplementary Table 22.** Linear mixed-effects model results from a distance model predicting neural pattern similarity that included the hippocampus and vmPFC. Source data are provided as a Source Data file. SE = standard error. No correction for multiple comparisons was applied.

| **Variable** | ***β*** | **SE** | ***t*** | ***p*** |
|---|---|---|---|---|
| (Intercept) | 0.036 | 0.037 | 0.982 | 0.326 |
| Day 2 | 0.002 | 0.014 | 0.138 | 0.890 |
| Day 3 | -0.003 | 0.012 | -0.205 | 0.837 |
| distance | 0.004 | 0.002 | 1.853 | 0.064 |
| region | -0.005 | 0.010 | -0.544 | 0.587 |
| Day 2 X distance | -0.003 | 0.003 | -1.025 | 0.306 |
| Day 3 X distance | -0.002 | 0.003 | -0.572 | 0.567 |
| Day 2 X region | 0.022 | 0.014 | 1.543 | 0.123 |
| Day 3 X region | -0.002 | 0.014 | -0.126 | 0.900 |

| | | | | |
|---|---|---|---|---|
| distance X region | -0.002 | 0.004 | -0.525 | 0.600 |
| Day 2 X distance X region | -0.003 | 0.005 | -0.663 | 0.508 |
| Day 3 X distance X region | 0.001 | 0.005 | 0.156 | 0.876 |

**Supplementary Table 23.** Linear mixed-effects model results from a distance model predicting neural pattern similarity that included the hippocampus, vmPFC, and a visual control region. The visual control region served as a baseline. Source data are provided as a Source Data file. SE = standard error. No correction for multiple comparisons was applied.

| **Variable** | ***β*** | **SE** | ***t*** | ***p*** |
|---|---|---|---|---|
| (Intercept) | 0.537 | 0.037 | 14.413 | **<2e-16** |
| Day 2 | -0.087 | 0.020 | -4.329 | **6.55e-5** |
| Day 3 | -0.059 | 0.022 | -2.712 | **0.009** |
| distance | -0.002 | 0.003 | -0.628 | 0.530 |
| region (hippocampus) | -0.506 | 0.011 | -46.889 | **<2e-16** |
| region (vmPFC) | -0.483 | 0.013 | -38.294 | **<2e-16** |
| Day 2 X distance | 0.008 | 0.005 | 1.655 | 0.098 |
| Day 3 X distance | -0.001 | 0.005 | -0.233 | 0.816 |
| Day 2 X region (hippocampus) | 0.077 | 0.015 | 5.102 | **3.38e-7** |
| Day 3 X region (hippocampus) | 0.047 | 0.015 | 3.079 | **0.002** |

| | | | | |
|---|---|---|---|---|
| Day 2 X region (vmPFC) | 0.089 | 0.017 | 5.129 | **2.92e-7** |
| Day 3 X region (vmPFC) | 0.028 | 0.018 | 1.549 | 0.121 |
| distance X region (hippocampus) | 0.006 | 0.004 | 1.387 | 0.165 |
| distance X region (vmPFC) | 0.005 | 0.005 | 1.073 | 0.283 |
| Day 2 X distance X region (hippocampus) | -0.010 | 0.006 | -1.821 | 0.069 |
| Day 3 X distance X region (hippocampus) | -0.000 | 0.006 | -0.041 | 0.967 |
| Day 2 X distance X region (vmPFC) | -0.015 | 0.007 | -2.296 | **0.022** |
| Day 3 X distance X region (vmPFC) | -0.001 | 0.007 | -0.086 | 0.931 |

**Supplementary Table 24.** Linear mixed-effects model results from a model of local (within-track) distance, predicting neural pattern similarity in vmPFC. Source data are provided as a Source Data file. SE = standard error. No correction for multiple comparisons was applied.

| **Variable** | ***β*** | **SE** | ***t*** | ***p*** |
|---|---|---|---|---|
| (Intercept) | 0.144 | 0.167 | 0.866 | 0.387 |
| Day 2 | -0.005 | 0.027 | -0.174 | 0.862 |
| Day 3 | -0.024 | 0.027 | -0.867 | 0.386 |
| distance (link distance 2 > link distance 1) | -0.005 | 0.020 | -0.234 | 0.815 |
| Day 2 X distance | 0.025 | 0.028 | 0.910 | 0.363 |

| | | | | |
|---|---|---|---|---|
| Day 3 X distance | 0.021 | 0.029 | 0.745 | 0.456 |

**Supplementary Table 25.** Linear mixed-effects model results from the model of global (across-track) distance, predicting neural pattern similarity in vmPFC. Source data are provided as a Source Data file. SE = standard error. No correction for multiple comparisons was applied.

| **Variable** | ***β*** | **SE** | ***t*** | ***p*** |
|---|---|---|---|---|
| (Intercept) | 0.055 | 0.119 | 0.462 | 0.644 |
| Day 2 | 0.043 | 0.044 | 0.993 | 0.321 |
| Day 3 | -0.001 | 0.046 | -0.012 | 0.990 |
| distance | 0.008 | 0.009 | 0.955 | 0.340 |
| Day 2 X distance | -0.013 | 0.013 | -1.071 | 0.284 |
| Day 3 X distance | -0.004 | 0.013 | -0.320 | 0.749 |

**Supplementary Table 26.** Linear mixed-effects model results in a model predicting correlations between similarity matrices for vmPFC, hippocampus, and a visual control region (calcarine). Source data are provided as a Source Data file. SE = standard error. No correction for multiple comparisons was applied.

| **Variable** | ***β*** | **SE** | ***t*** | ***p*** |
|---|---|---|---|---|
| (Intercept) | 0.114 | 0.037 | 3.119 | **0.002** |
| scan session | -0.007 | 0.017 | -0.418 | 0.676 |
| comparison region | 0.167 | 0.052 | 3.233 | **0.002** |

| | | | | |
|---|---|---|---|---|
| scan session X comparison region | -0.022 | 0.024 | -0.911 | 0.364 |

**Supplementary Table 27.** Linear mixed-effects model results in a model of local (within-track) distance, predicting neural pattern similarity in an 8mm spherical vmPFC ROI. Source data are provided as a Source Data file. SE = standard error. No correction for multiple comparisons was applied.

| **Variable** | ***β*** | **SE** | ***t*** | ***p*** |
|---|---|---|---|---|
| (Intercept) | 0.031 | 0.023 | 1.354 | 0.176 |
| Day 2 | 0.010 | 0.035 | 0.293 | 0.769 |
| Day 3 | -0.027 | 0.031 | -0.845 | 0.398 |
| distance (link distance 2 > link distance 1) | -0.011 | 0.022 | -0.532 | 0.595 |
| Day 2 X distance | -0.020 | 0.031 | -0.669 | 0.503 |
| Day 3 X distance | 0.032 | 0.031 | 1.010 | 0.313 |

**Supplementary Table 28.** Linear mixed-effects model results in a model of global (across-track) distance, predicting neural pattern similarity in an 8mm spherical vmPFC ROI. Source data are provided as a Source Data file. SE = standard error. No correction for multiple comparisons was applied.

| **Variable** | ***β*** | **SE** | ***t*** | ***p*** |
|---|---|---|---|---|
| (Intercept) | 0.221 | 0.130 | 1.705 | 0.088 |
| Day 2 | 0.024 | 0.052 | 0.453 | 0.651 |

| | | | | |
|---|---|---|---|---|
| Day 3 | -0.033 | 0.050 | -0.660 | 0.509 |
| distance | -0.003 | 0.010 | -0.273 | 0.785 |
| Day 2 X distance | -0.013 | 0.014 | -0.955 | 0.339 |
| Day 3 X distance | 0.004 | 0.014 | 0.251 | 0.802 |

**Supplementary Table 29.** Linear mixed-effects model results for a context model predicting neural pattern similarity, fit to data from an 8mm spherical ROI in vmPFC. Source data are provided as a Source Data file. SE = standard error. No correction for multiple comparisons was applied.

| **Variable** | ***β*** | **SE** | ***t*** | ***p*** |
|---|---|---|---|---|
| (Intercept) | 0.089 | 0.038 | 2.346 | **0.019** |
| Day 2 | -0.023 | 0.022 | -1.032 | 0.302 |
| Day 3 | -0.024 | 0.016 | -1.527 | 0.127 |
| stimulus type (Landmark > Fractal) | -0.002 | 0.005 | -0.442 | 0.658 |
| context (same track > different tracks) | 0.005 | 0.005 | 1.097 | 0.273 |
| Day 2 X stimulus type | 0.009 | 0.007 | 1.295 | 0.195 |
| Day 3 X stimulus type | 0.022 | 0.007 | 3.144 | **0.002** |
| Day 2 X context | -0.013 | 0.007 | -1.954 | 0.051 |
| Day 3 X context | -0.007 | 0.007 | -1.031 | 0.303 |

| | | | | |
|---|---|---|---|---|
| stimulus type X context | 0.012 | 0.010 | 1.206 | 0.228 |
| Day 2 X stimulus type X context | -0.011 | 0.014 | -0.797 | 0.426 |
| Day 3 X stimulus type X context | -0.020 | 0.014 | -1.407 | 0.159 |

**Supplementary Table 30.** Linear mixed-effects model results from a context model predicting neural pattern similarity that included the hippocampus, EC, and a visual control region. The visual control region served as a baseline. Source data are provided as a Source Data file. SE = standard error. No correction for multiple comparisons was applied.

| **Variable** | ***β*** | **SE** | ***t*** | ***p*** |
|---|---|---|---|---|
| (Intercept) | 0.547 | 0.020 | 26.981 | **<2e-16** |
| Day 2 | -0.029 | 0.014 | -2.035 | **0.042** |
| Day 3 | -0.024 | 0.018 | -1.346 | 0.178 |
| distance | 0.002 | 0.004 | 0.430 | 0.667 |
| region (hippocampus) | -0.440 | 0.003 | -148.995 | **<2e-16** |
| region (EC) | -0.458 | 0.003 | -156.298 | **<2e-16** |
| Day 2 X distance | -0.003 | 0.006 | -0.467 | 0.640 |
| Day 3 X distance | -0.004 | 0.006 | -0.596 | 0.551 |
| Day 2 X region (hippocampus) | 0.006 | 0.004 | 1.521 | 0.128 |
| Day 3 X region (hippocampus) | 0.009 | 0.004 | 2.202 | **0.028** |

| | | | | |
|---|---|---|---|---|
| Day 2 X region (EC) | -0.008 | 0.004 | -2.019 | **0.043** |
| Day 3 X region (EC) | 0.017 | 0.004 | 4.354 | **1.34e-5** |
| distance X region (hippocampus) | -0.001 | 0.005 | -0.114 | 0.909 |
| distance X region (EC) | 0.003 | 0.005 | 0.579 | 0.563 |
| Day 2 X distance X region (hippocampus) | -0.001 | 0.007 | -0.119 | 0.905 |
| Day 3 X distance X region (hippocampus) | 0.001 | 0.008 | 0.185 | 0.853 |
| Day 2 X distance X region (EC) | -0.010 | 0.007 | -1.370 | 0.171 |
| Day 3 X distance X region (EC) | -0.006 | 0.008 | -0.792 | 0.429 |

**Supplementary Table 31.** Linear mixed-effects model results from a context model predicting neural pattern similarity, fit to data from a visual control region. Source data are provided as a Source Data file. SE = standard error. No correction for multiple comparisons was applied.

| **Variable** | ***β*** | **SE** | ***t*** | ***p*** |
|---|---|---|---|---|
| (Intercept) | 0.554 | 0.043 | 13.030 | **<2e-16** |
| Day 2 | -0.021 | 0.025 | -0.820 | 0.412 |
| Day 3 | -0.039 | 0.041 | -0.944 | 0.345 |
| stimulus type (landmark > fractal) | -0.024 | 0.003 | -7.009 | **2.45e-12** |
| context (same track > different tracks) | 0.001 | 0.003 | 0.245 | 0.806 |

| | | | | |
|---|---|---|---|---|
| Day 2 X stimulus type | 0.005 | 0.005 | 1.088 | 0.277 |
| Day 3 X stimulus type | -0.002 | 0.005 | -0.407 | 0.684 |
| Day 2 X context | -0.003 | 0.005 | -0.717 | 0.474 |
| Day 3 X context | -0.002 | 0.005 | -0.416 | 0.678 |
| stimulus type X context | -0.008 | 0.007 | -1.252 | 0.211 |
| Day 2 X stimulus type X context | -0.004 | 0.009 | -0.377 | 0.706 |
| Day 3 X stimulus type X context | 0.014 | 0.010 | 1.509 | 0.131 |

**Supplementary Table 32.** Linear mixed-effects model results from a model of local (within-track) distance predicting neural pattern similarity, fit to data from a visual control region. Source data are provided as a Source Data file. SE = standard error. No correction for multiple comparisons was applied.

| **Variable** | ***β*** | **SE** | ***t*** | ***p*** |
|---|---|---|---|---|
| (Intercept) | 0.317 | 0.135 | 2.343 | **0.019** |
| Day 2 | -0.040 | 0.030 | -1.351 | 0.177 |
| Day 3 | -0.025 | 0.044 | -0.577 | 0.564 |
| distance (link distance 2 > link distance 1) | 0.014 | 0.016 | 0.870 | 0.384 |
| Day 2 X distance | 0.007 | 0.022 | 0.304 | 0.761 |
| Day 3 X distance | -0.035 | 0.022 | -1.570 | 0.117 |

**Supplementary Table 33.** Linear mixed-effects model results from a model of global (across-track) distance predicting neural pattern similarity, fit to data from a visual control region. Source data are provided as a Source Data file. SE = standard error. No correction for multiple comparisons was applied.

| **Variable** | ***β*** | **SE** | ***t*** | ***p*** |
|---|---|---|---|---|
| (Intercept) | 0.418 | 0.095 | 4.379 | **1.31e-5** |
| Day 2 | -0.003 | 0.040 | -0.082 | 0.935 |
| Day 3 | -0.061 | 0.051 | -1.190 | 0.234 |
| distance | -0.005 | 0.007 | -0.783 | 0.434 |
| Day 2 X distance | -0.003 | 0.010 | -0.261 | 0.794 |
| Day 3 X distance | 0.007 | 0.010 | 0.727 | 0.468 |
